# A Bayesian Heteroscedastic GLM with Application to fMRI Data with Motion Spikes

**DOI:** 10.1101/091066

**Authors:** Anders Eklund, Martin A. Lindquist, Mattias Villani

## Abstract

We propose a voxel-wise general linear model with autoregressive noise and heteroscedastic noise innovations (GLMH) for analyzing functional magnetic resonance imaging (fMRI) data. The model is analyzed from a Bayesian perspective and has the benefit of automatically down-weighting time points close to motion spikes in a data-driven manner. We develop a highly efficient Markov Chain Monte Carlo (MCMC) algorithm that allows for Bayesian variable selection among the regressors to model both the mean (i.e., the design matrix) and variance. This makes it possible to include a broad range of explanatory variables in both the mean and variance (e.g., time trends, activation stimuli, head motion parameters and their temporal derivatives), and to compute the posterior probability of inclusion from the MCMC output. Variable selection is also applied to the lags in the autoregressive noise process, making it possible to infer the lag order from the data simultaneously with all other model parameters. We use both simulated data and real fMRI data from OpenfMRI to illustrate the importance of proper modeling of heteroscedasticity in fMRI data analysis. Our results show that the GLMH tends to detect more brain activity, compared to its homoscedastic counterpart, by allowing the variance to change over time depending on the degree of head motion.

## 1. Introduction

Functional magnetic resonance imaging (fMRI) is a non-invasive technique that has become the de facto standard for imaging human brain function in both healthy and diseased populations. The standard approach for analyzing fMRI data is to use the general linear model (GLM), proposed by Friston et al. [12]. The standard GLM has been extremely successful in a large number of empirical studies, but relies on a number of assumptions, including linearity, independency, Gaussianity and homoscedasticity (constant variance). Much work has been done to relax the assumption of independent errors, and several alternative noise models have been proposed [11, 41, 21, 19, 7]. In addition, it has also been investigated whether results are improved by using a Rician noise model [16, 1, 23, 35], instead of a Gaussian. While heteroscedastic models exist for group analyses [2, 39, 5], the homoscedasticity assumption for single subject analysis has received little attention. Luo and Nichols [22] used the Cook-Weisberg test for homoscedasticity to detect problematic voxels, but did not propose a heteroscedastic model to handle these. Diedrichsen and Shadmehr [6] claim that the homoscedasticity assumption is often violated in practice due to head motion, and propose an algorithm that estimates the noise variance separately at each time point. The estimated variances are then used to perform weighted least squares regression. The aim of this study is to further explore the appropriateness of the homoscedasticity assumption for single subject fMRI analysis, and evalute the effects of deviations from it.

### 1.1. Is fMRI noise heteroscedastic?

Consider a simple simulation where actual head motion is applied to a single volume from a real fMRI dataset, to generate a new 4D fMRI dataset where all the signal variation comes from simulated motion. For each time point, the corresponding head motion parameters are used to translate and rotate the first volume in the dataset (using interpolation), and the transformed volume is saved as the volume for that specific time point. Even if motion correction is applied to the simulated dataset, the dataset will still contain motion related signal variation [15], due to the fact that the interpolation mixes voxels with low and high signal intensity (especially at the edge of the brain, and at the border between different tissue types). It is therefore common to include the estimated head motion parameters in the design matrix, to regress out any motion related variance that remains after the motion correction, and to also account for spin-history artifacts. It is also common to include the temporal derivative of the head motion parameters, to better model motion spikes. Figure 1 shows a single time series from an fMRI dataset with simulated motion, before and after motion correction, and one of the head motion parameters. The selected voxel is at the border between white and gray matter. Figure 2 shows three residual time series calculated using three different design matrices (and ordinary least squares regression), the first containing only an intercept and time trends, the second also containing motion covariates, and the third also containing the temporal derivative of the head motion. It is clear that using the estimated head motion as additional covariates removes most of the motion related variance, but not all of it. The residual time series still contain effects of a motion spike, which makes the noise heteroscedastic.

**Figure 1:**
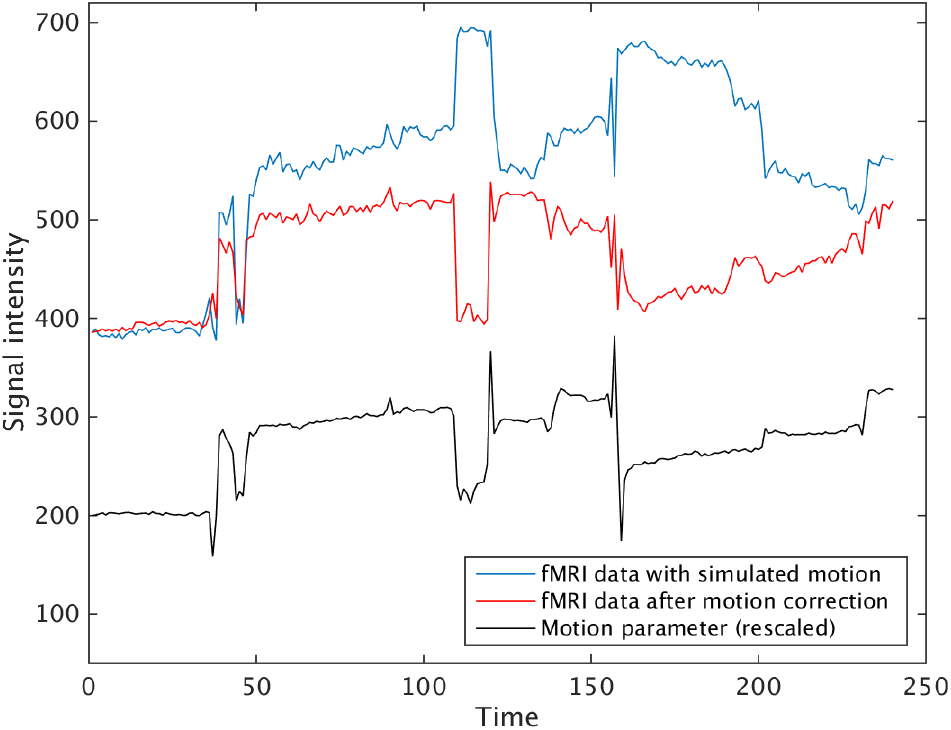
A single time series from a simulated fMRI dataset, before and after motion correction. All the signal variation comes from simulated motion, and is due to interpolation artefacts. Note that the motion corrected data still has a high signal variance, which is correlated with the head motion.

**Figure 2:**
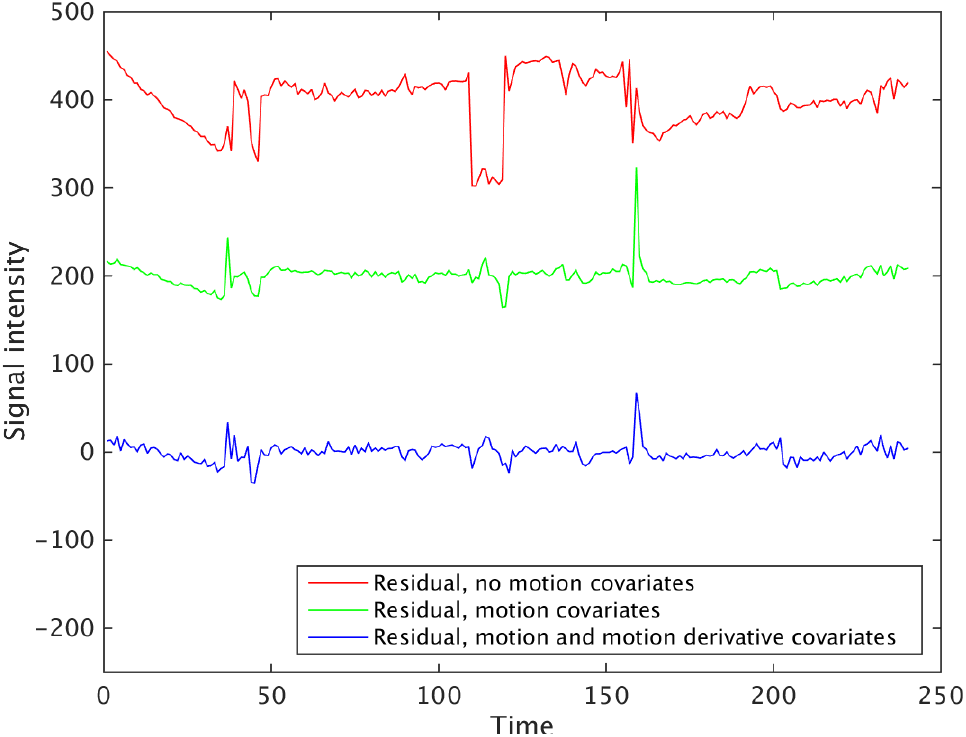
Residual time series obtained after fitting models with three different design matrices. The first design matrix only contains covariates for the intercept and time trends (4 covariates in total). The second design matrix also contains head motion covariates (10 covariates in total), and the third design matrix also contains the temporal derivative of the head motion (16 covariates in total). For visualization purposes, a mean of 200 was added to the green residual, and a mean of 400 was added to the red residual. Note that a motion spike is still present in the green and the blue residual, making the noise heteroscedastic.

It should also be stressed that real fMRI data are far more complicated, for example due to the fact that each fMRI volume is sampled one slice at a time. Another problem is so-called ’spin-history’ effects which alter the signal intensity of volumes following the motion spike, because the head motion changes the excitation state of the spins of the protons (thereby interrupting the steady state equilibrium). For this reason, a number of volumes after the motion spike should also be downweighted, and not only volumes during the motion spike.

### 1.2. Modeling the heteroscedasticity

We propose a Bayesian heteroscedastic extension of the GLM, which uses covariates for both the mean and variance, and also incorporates an autoregressive noise model. We develop highly efficient Markov Chain Monte Carlo (MCMC) algorithms for simulating from the joint posterior distribution of all model parameters. Allowing for heteroscedasticity, where the noise variance is allowed to change over time, has the effect of automatically discounting scans with large uncertainty when inferring brain activity or connectivity. One way of thinking of this effect is in terms of weighted least squares estimation, where the optimal weights are learned from the data.

### 1.3. Is fMRI noise heteroscedastic in all voxels?

Figure 3 shows three residual time series for a voxel in gray matter (close to the voxel shown in Figure 2). Clearly, this voxel has a very low correlation with the simulated motion, and the residuals are not heteroscedastic. It is therefore not optimal to use the same weights in all voxels. Compared to the work by Diedrichsen and Shadmehr [6], our Bayesian approach independently estimates a heteroscedastic model for each voxel, instead of using variance scaling parameters that are the same for all voxels. Furthermore, Diedrichsen and Shadmehr [6] used a fix autoregressive (AR) model for the noise (AR(1) + white noise with the AR parameter fixed to 0.2, as in the SPM software package), while we estimate an AR(k) model in each voxel. The fixed AR(1) model used by SPM has been shown to perform poorly [7], especially for short repetition times made possible with recently developed MR scanner sequences.

**Figure 3:**
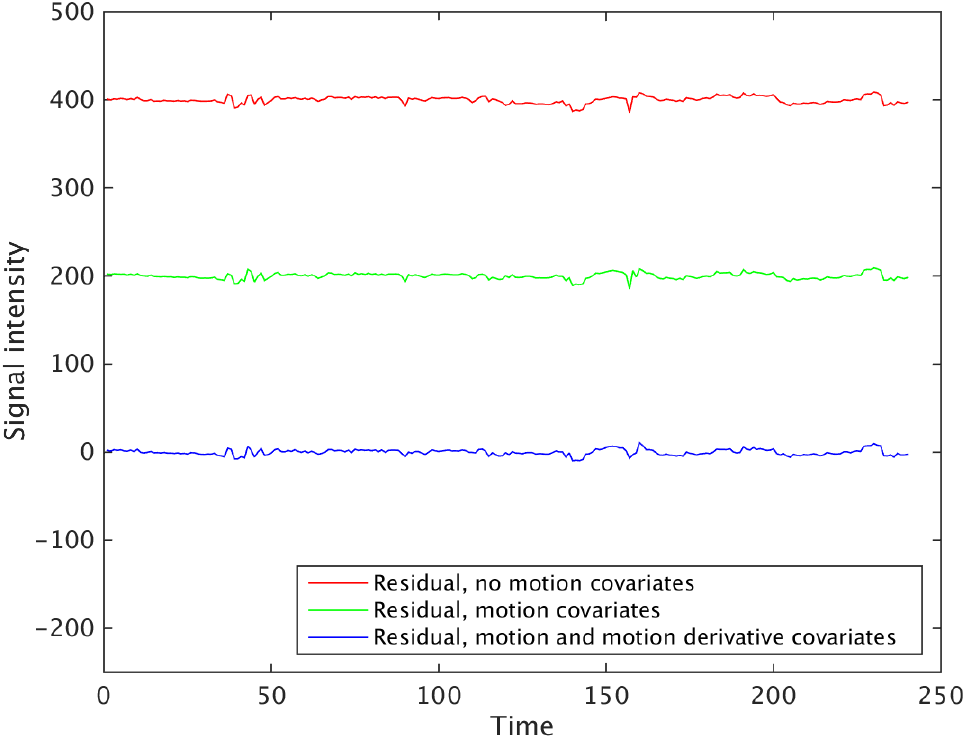
Residual time series obtained after fitting models with three different design matrices. The first design matrix only contains covariates for intercept and time trends (4 covariates in total). The second design matrix also contains head motion covariates (10 covariates in total), and the third design matrix also contains the temporal derivative of the head motion (16 covariates in total). For visualization purposes, a mean of 200 was added to the green residual, and a mean of 400 was added to the red residual. Note that the time series in this voxel has a very low correlation with the simulated motion, and the residuals are therefore homoscedastic.

Our Bayesian approach also differs from recently developed methods used in the field, where scrubbing or censoring is used to remove volumes with excessive head motion [30, 28, 32]. Such approaches are ad hoc in the sense that an arbitrary motion threshold first needs to be applied, to determine which volumes to remove or censor. Another problem with these approaches is that they can significantly alter the temporal structure of the data.

### 1.4. Variable selection

It can be difficult to determine which variables to include in the design matrix (i.e., the mean function) of the GLM, including those that capture scanner drift, or residual head movement effects after motion correction. It can be even more difficult to choose the appropriate explanatory variables to use in the variance function. For this reason we introduce variable selection priors in both the mean and variance function, which has the effect of automatically determining the set of explanatory variables; more precisely, we obtain the posterior inclusion probability for each of the candidate variables and the posterior distribution of their effect sizes from a single MCMC run. In addition, we have a third variable selection prior acting on the lags of the AR noise process which allows us to estimate the model order of the AR process directly from the data. This aspect is particularly important for high (sub-second) temporal resolution data. Our analysis here is massively univariate without modeling spatial dependencies, however we discuss possible extensions to spatial models in the Discussion.

## 2. GLM with heteroscedastic autoregressive noise

We propose the following voxel-wise GLM with heteroscedastic noise innovations (GLMH) for blood oxygenation level dependent (BOLD) time series:

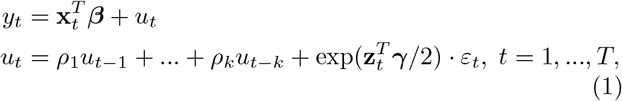

where *y_t_* is the observed fMRI signal at time *t*, x_*t*_ is a vector with *p* covariates for modeling the mean, *ε_t_* is zero mean Gaussian white noise with unit variance, and z_*t*_ is a vector of *q* covariates for modeling the variance of the heteroscedastic noise innovations as ln 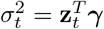. The logarithm of the variance is modelled as a linear regression, to enable unrestricted estimation of *γ* while still guaranteeing a positive variance. Note that we are here using the logarithmic link function for the variance, but our methodology is applicable to any invertible and twice-differentiable link function. The GLMH model introduces heteroscedasticity through noise innovations with the effect that a large variance at time *t* is likely to generate a large innovation in the *u_t_* equation, which is propagated through the autoregressive structure. The effect is that the noise remains large in subsequent scans, which is desireable as it has been shown that motion related signal changes can persist more than 10 seconds after motion ceases [28] (for example due to spin-history effects, as mentioned in the Introduction).

Let **y** = (*y_1_,…,y_T_*)^*T*^ be a *T*-dimensional vector consisting of observed fMRI signals at a specific voxel and define *u* and *ε* analogously. Also, define **X** = (**x**_1_,…, **x**_*T*_)^*T*^ and **Z** = (**z**_1_,…, **z**_*T*_)^*T*^ to be *T* × *p* and *T* × *q* matrices consisting of covariates. Further, let ***ρ*** = (*ρ*_1_, …, *ρ_k_*)^*T*^. The GLMH model can then be written as follows:

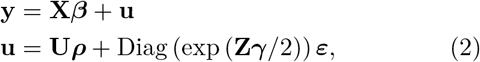

where **U** is a *T* × *k* matrix consisting of lagged values of **u**, assuming that *k* pre-sample observations are available. Figure 4 shows an example of **X** and **Z** for a subject with several motion spikes.

**Figure 4:**
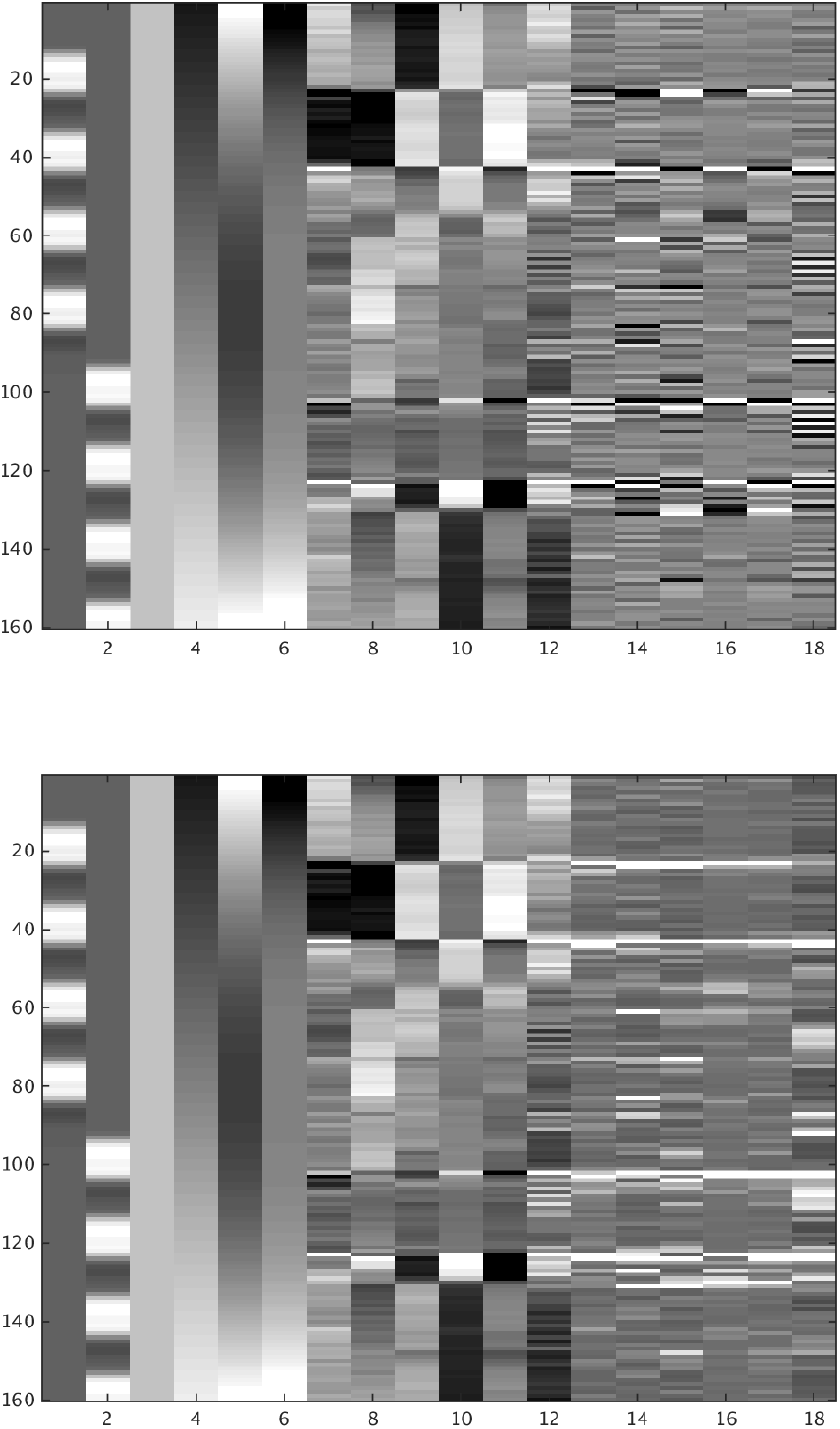
An example of **X** (top) and **Z** (bottom) for a subject with several motion spikes. The data consists of 160 time points, and 18 covariates are here used to model both the mean (**X**) and the variance (**Z**). The first two covariates (from the left) represent two different tasks, the following four covariates represent intercept, linear trend, quadratic trend and cubic trend, the following six covariates represent the estimated head motion, and the last six covariates represent the temporal derivative of the head motion. The only difference between **X** and **Z** is that the absolute value of the temporal derivative is used for **Z**, as a motion spike should always lead to an increase of the variance. All covariates (except the intercept) are standardized to have zero mean and unit variance, which often leads to a better convergence of the MCMC chain.

## 3. Bayesian inference

We begin by defining the binary indicators ℐ_*β*_, ℐ_*γ*_, and ℐ_*ρ*_, which are used for variable selection purposes. Here ℐ_*β*_ is a *p* × 1 vector whose jth element takes the value 1 if _*j*_ is non-zero and 0 otherwise. The indicators ℐ_*γ*_ and ℐ_*ρ*_ are defined analogously. We take a Bayesian approach with the aim of computing the joint posterior distribution *p*(***β***, ***γ***, ***ρ***, ℐ_*β*_, ℐ_*γ*_, ℐ_*ρ*_|**y**, **X**, **Z**). This distribution is intractable and we use Metropolis-within-Gibbs sampling [4] to generate draws from the joint posterior. The algorithm iterates between the following three full conditional posteriors:

1. (*β*, ℐ_β_)|**y, X, Z,.**
2. (*ρ,* ℐ_*ρ*_) **y, X, Z,.**
3. (*γ,* ℐ*γ*) **y, X, Z,.**

where · denotes all other model parameters.

### 3.1. Prior distribution

We assume prior independence between ***β***, ***γ*** and ***ρ***, and let

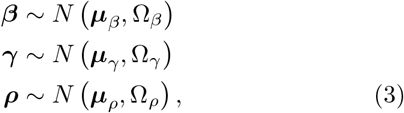

where 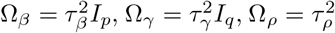 Diag 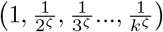 and ***μ***_*ρ*_ = (*r*, 0,…, 0)^*T*^. The prior mean ***μ**_β_* is set to 0 for all parameters, except for the term corresponding to the intercept which is set to 800. The prior mean ***μ**_γ_* is set equal to 0 for all parameters. Note that the *N*(***μ**_ρ_*, Ω_*ρ*_) prior centers the prior on the *AR*(1) process *u_t_* = *r* · *u*_*t*-1_ + *ε_t_*, with coefficients corresponding to longer lags more tightly shrunk toward zero. We also restrict the prior on ***ρ*** to the stationarity region. The user is required to specify the prior hyperparameters *τ_β_, τ_*γ*_, τ_ρ_, r* and *ζ*. As default values we use *τ_β_* = *τ_γ_* = 10, *τ_ρ_* = 1, *r* = 0.5 and *ζ* = 1, providing a rather uninformative prior. A more complex prior, which for example allows for prior dependence between *β* and *γ*, can easily be incorporated into our framework.

### 3.2. Variable selection

Our MCMC algorithm presented in Section 3.4 performs Bayesian variable selection among both sets of covariates, x_*t*_ (mean) and z_*t*_ (variance), using a spike and slab prior [13, 18]. We also use Bayesian variable selection in the AR noise process, thereby automatically learning about the order *k* of the AR process. The first element of *β* and *γ* (i.e., the intercepts in the mean and log variance, respectively) are not subject to variable selection. To describe the variable selection prior, let us focus on *β*. Let *β_ℐβ_* denote the subset of regression coefficients selected by ℐ*β*. To allow for variable selection we take the prior for the unrestricted ***β*** ~ *N*(***μ**_β_*, **Ω**_*β*_) and condition on the zeros dictated by ℐ_*β*_. Since all our prior covariance matrices are diagonal, the conditional distributions are simply the marginal distributions, e.g. 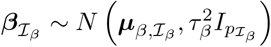, where ***μ**_β, ℐβ_* is the subset of elements of ***μ*** corresponding to ℐ_*β*_, and *p_ℐβ_* is the number of elements in 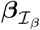. To complete the variable selection prior we let the elements of ℐ_*β*_ be apriori independent and Bernoulli distributed with Pr(*I*_*β,j*_ = 1) = *π_β_*. The default values for *π_β_* and *π_γ_* are 0.5. The default value for *π_ρ_* is 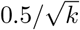 for lag *k*, giving 0.5, 0.35, 0.29 and 0.25 for an AR(4) process. We also experiment with a hierarchical prior where the n are assigned Beta priors, see below. The extension to a spatial prior on the variable selection indicators is also discussed below.

### 3.3. Variable selection in linear regression using MCMC

This section describes how to simulate from the joint posterior of the regression coefficients, and their variable selection indicators in the Gaussian linear regression model with unit noise variance

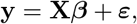

where *ε* = (*ε*_1_, *…, ε_T_*)^*T*^ and 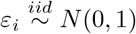 This will be an important building block in our Metropolis-within-Gibbs algorithms described in Sections 2 and 3.4. Similar to Smith and Kohn [34], we sample *β* jointly with its variable selection indicators ℐ (we drop the subscript *β* here) by first generating from the marginal posterior *p*(ℐ|**y, X**) followed by a draw from *p*(*β*| ℐ, **y, X**). A draw from *p*(*β*| ℐ, **y, X**) is easily obtained by sampling the non-zero elements of *β* as

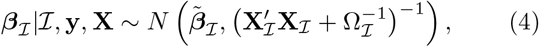

where **X**_ℐ_ is the *T* × _*pℐ*_ matrix with covariates from the subset ℐ, Ω_ℐ_ is the prior covariance for *β*_ℐ_ | ℐand *β̃*_ℐ_ is given in Appendix B. A closed form expression for *p*(ℐ|**y, X**), the marginal posterior of ℐ, is given in Appendix B, from which we can obtain *p*(ℐ_*j*_ |**y, X**, ℐ_−*j*_) *∞ p*(ℐ|**y, X**), where *ℐ_-j_* denotes ℐ with the *j*th element excluded. Simulating from the joint posterior of *β* and ℐ is therefore acheived by simulating from each *p*(ℐ_*j*_ |**y, X**, ℐ_-*j*_) in turn, followed by sampling of *β_ℐ_* from (4).

### 3.4. MCMC for the GLMH model

#### Updating (*β, ℐ _β_*)

To sample from the full conditional posterior of (*β*, ℐ _*β*_) conditional on ***ρ*** and *γ*, let us re-formulate the model as

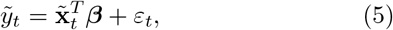

where 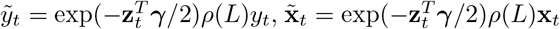 and *ρ*(*L*) = 1 — *ρ*_1_*L* — … — *ρ*_*k*_*L*^*k*^ is the usual lag polynomial in the lag operator *L^k^_yt_* = *y_t-k_* from time series analysis. The Jacobian of the transformation 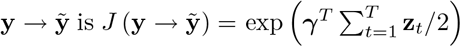, which can be seen as follows. The inverse transformation is 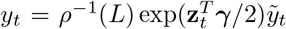, where *ρ*^-1^(*L*) = 1 + *ψ*_1_*L* + *ψ*_2_*L*^2^ + … is the inverse lag polynomial for some coefficients *ψ*_1_, *ψ*_2_,…. This system of equations is recursive so the Jacobian is 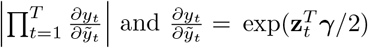 which proves the result. Note that *J*(**y** → **ỹ**) does not depend on *β* and can therefore be ignored when deriving the full conditional posterior of *β*. Now, *β* in (5) are the coefficients in a linear regression with unit noise variance and we can draw from the full conditional *p*(*β*, ℐ _*β*_|**y, X, Z**, ·) as described in Section 3.3 with y and **X** replaced by ỹ and X̃, respectively.

#### Updating (*ρ, ℐ_ρ_*)

The AR process can be rewritten as

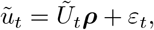

where 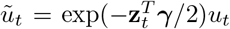. The Jacobian of this transformation is 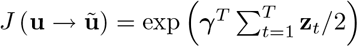 which does not depend on ***ρ*** and can therefore be ignored when updating ***ρ***. Now, ***ρ*** are the coefficients in a linear regression with unit noise variance and we can draw from the full conditional *p*(***ρ***, ℐ _*ρ*_ |**y, X, Z**, ·) as described in Section (3.3).

#### Updating (*γ*, ℐ *_ρ_*)

The full conditional posterior of (*γ*, *ℐ _γ_*) is a complicated distribution which we can not easily sample from. However, it is clear from the model

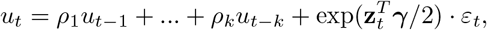

that the conditional likelihood of *γ* is of the form described in Villani et al. [38] where the observations (the *u_t_* in this case) are conditionally independent and *γ* enters each factor in the likelihood linearly 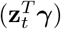 through a scalar valued quantity 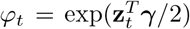. The MCMC update with a finite step Newton proposal with variable selection described in [37, 38] can therefore be used. In fact, Villani et al. [37] contains the details for the Gaussian heteroscedastic regression, which is exactly the model when we condition on *β* (since **u** is then known). The algorithm in [38] proposes *γ* and *ℐ _γ_* jointly by randomly changing a subset of the indicators in *ℐ _γ_* followed by a proposal from *γ* | ℐ _*γ*_ using a multivariate-*t* distribution tailored to the full conditional posterior. The tailoring is acheived by taking a small number of Newton steps toward the posterior mode, and using the negative inverse Hessian at the terminal point as the covariance matrix in the multivariate-*t* proposal distribution. The update is fast, since the Jacobian and Hessian can be computed in closed form using the chain rule and compact matrix algebra. It is also possible to compute the expected Hessian (Fisher information) in closed form. The expected Hessian tends to be more stable numerically with only marginally worse tailoring to the posterior. Note also that the Newton iterations always start from the current value of *γ*, which is typically not far from the mode, so even one or two Newton steps are usually sufficient. We refer to Villani et al. [38] for details of the general algorithm, and to Villani et al. [37] for expressions of the Jacobian, Hessian and expected Hessian for *γ.*

#### Updating π_β_ and π_γ_

The inclusion probabilities *π_β_* and *π_γ_* for the variable selection can also be updated in every MCMC iteration[18] (updating *π_ρ_* is in principle straightforward, but there is very little information about *π_ρ_*, due to the low number of AR parameters). Let the prior for *π_β_* and *π_γ_* be Beta(*a*, *b*). The conditional posterior for *π_β_* is then given by Beta 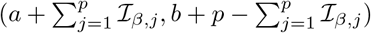, where *p* is the number of covariates and *ℐ _β,j_* is the binary indicator varible for covariate *j*. The posterior for *π_γ_* is defined analogously. We use *a* = *b* = 3 which gives a prior with a mean of 0.5. The complete algorithm becomes

1. (*β*, *ℐ _β_*)|**y,X,Z, ·**
2. (*π_β_*)|**y, X, Z**, ·
3. (***ρ***, ℐ _*ρ*_)|**y, X, Z**, ·
4. (*γ*, ℐ _*γ*_)|**y, X, Z**, ·
5. (*π_γ_*)|**y, X, Z**, ·

where · denotes all other model parameters.

#### Spatial variable selection prior

Since ***β*** and ***ρ*** both appear as coefficients in linear regressions (conditional on the other parameters), it is straightforward to extend our variable selection for *ℐ _β_* and *ℐ _ρ_* to have a spatial binary Markov random field prior following Smith and Fahrmeir [33], but would naturally add to the processing time. A spatial prior on ℐ _*γ*_ is more difficult since *γ* does not appear linearly in the model, even conditional on the other parameters, and can therefore not be integrated out analytically as in Smith and Fahrmeir [33]. We leave such an extension to future work.

## 4. Implementation

A drawback of using MCMC is that processing of a single fMRI dataset can take several hours [40, 31]. Our implementation of the heteroscedastic GLM is therefore written in C++, using the Eigen library [17] for all matrix operations. The random number generators available in the C++ standard library (available from C++ 2011) were used, together with the Eigen library, to make random draws from multivariate distributions. The OpenMP (Open Multi Processing) library was used to take advantage of all the CPU cores, by analyzing several voxels in parallel. For all analyses the number of Newton steps is set to 2. To lower the processing time, the variable selection indicators for the variance covariates are only updated in 60% of the draws. See Appendix A for more information about the implementation. The code is available at https://github.com/wanderine/HeteroscedasticfMRI

## 5. Results

### 5.1 Simulated data

#### 5.1.1. GLMH vs Bayesian GLM with homoscedastic noise

To verify that the heteroscedastic model works as expected, and to compare it to a homoscedastic model for data with a known activity pattern, the algorithms were applied to simulated data with homoscedastic and heteroscedastic noise. The simulated data were created using (posterior mean) beta estimates from spatially smoothed real fMRI data (with several motion spikes), together with the applied design matrix, to create a timeseries in each voxel. The design matrix consisted of an intercept, time trends for modeling drift (linear, quadratic and cubic), activity covariates, estimated head motion parameters and their temporal derivative (in total 16 covariates in addition to the activity covariates, see Figure 4 for an example). The simulated data thereby contain spatial correlation as well as correlation between the covariates. Beta values for active voxels were generated from a *abs(N*(0, 9)) + 3 distribution, and for non-active voxels from a *N*(0,0.06) distribution. The simulated activity is thereby very easy to detect, and the difficult part is to model the heteroscedastic noise.

For approximately half of the active voxels, heteroscedastic noise was added according to Equation 1. For one covariate at a time (either an activation or head motion covariate), the corresponding *γ* parameter was set to 1, 2, or 3. For one covariate representing the (absolute value of the) temporal derivative of the head motion, the *γ* parameter was instead set to 1, 1.25 or 1.5 (as motion spikes can be rather large, and thereby make the simulation unrealistic). To simulate simultaneous heteroscedasticity from several covariates, the *γ* parameters for the activity and the head motion covariates were simultaneously set to 1, 2, or 3, while the *γ* parameter for the derivated head motion covariate was set to 1.25 for all cases. The *γ* parameter for the intercept covariate was always set to 1, and all other *γ* parameters were set to 0. For all other voxels, homoscedastic noise was added (*γ* =1 for the intercept only). The four autocorrelation parameters were set to 0.4, 0.2, 0.1 and 0.05, respectively. The simulated data thereby consists of four regions; active voxels with homoscedastic or heteroscedastic noise, and non-active voxels with homoscedastic or heteroscedastic noise. To lower the processing time, only a single slice of data was simulated. See Figure 5 for the gray matter mask, the mask for active voxels and the mask for voxels with heteroscedastic noise. Figure 6 shows one simulated time series with homoscedastic noise, and two simulated time series with heteroscedastic noise.

**Figure 5:**
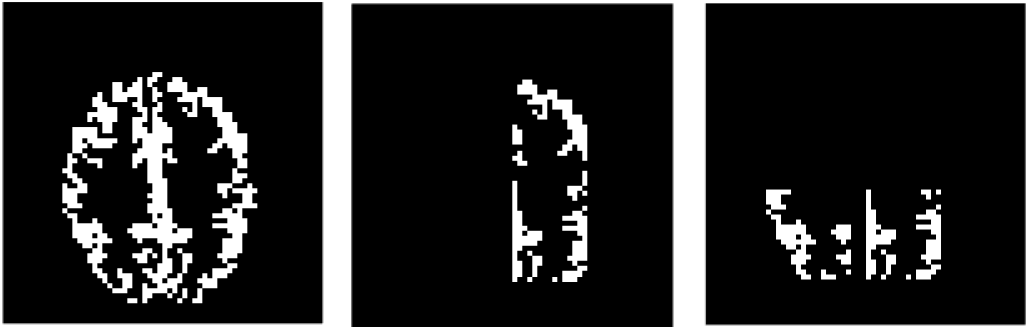
Left: A mask for gray matter voxels. Middle: Voxels with simulated activity. Right: Voxels with heteroscedastic noise. The simulated data consists of four regions; active voxels with homoscedastic or heteroscedastic noise, and non-active voxels with homoscedastic or heteroscedastic noise.

**Figure 6:**
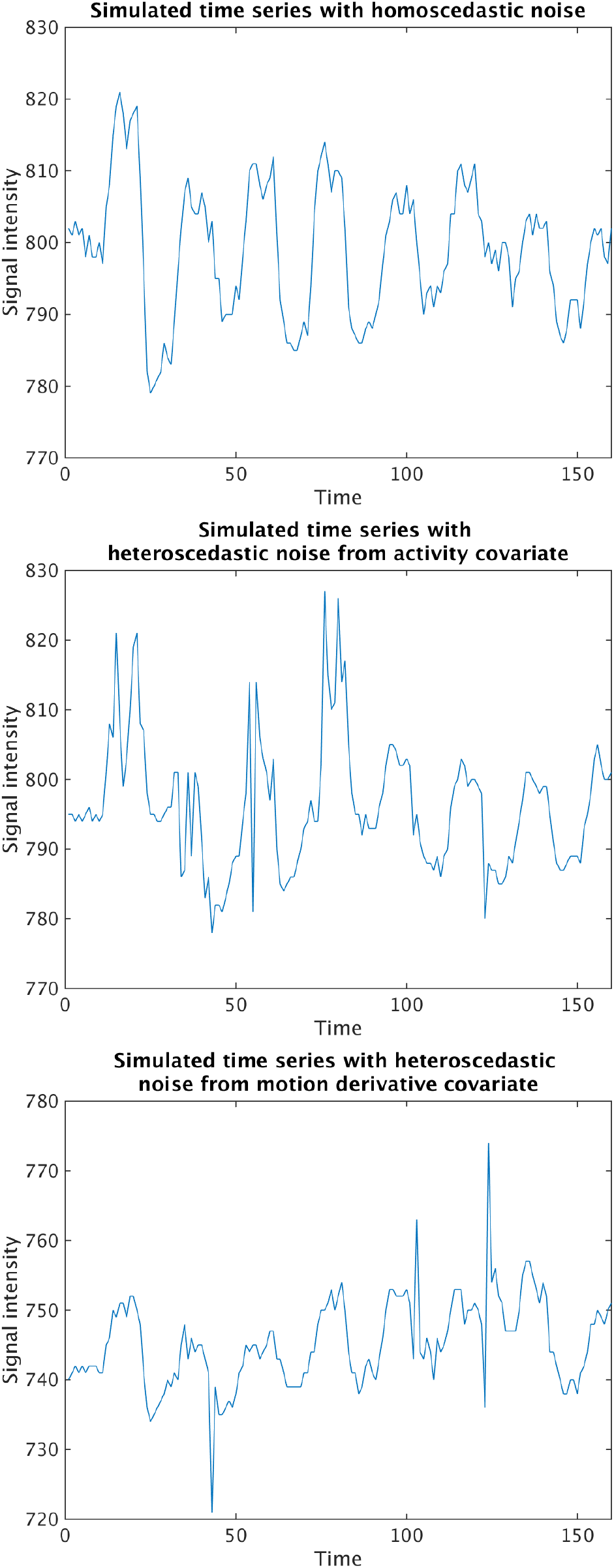
Top: A simulated time series with homoscedastic noise. Middle: A simulated time series with heteroscedastic noise, where the variance is modelled as a function of an activity covariate (the variance is higher for the first part of the dataset, which corresponds to the first activity covariate). Bottom: A simulated time series with heteroscedastic noise, where the variance is modelled as a function of a covariate representing the temporal derivative of the head motion. In all cases the simulated brain activity is rather strong, but the heteroscedastic noise makes it difficult to detect activity using homoscedastic methods.

For each simulated dataset, the analysis was performed (i) including only an intercept for the variance (i.e., a homoscedastic model) and (ii) including all covariates for the variance (i.e., a heteroscedastic model). In both cases all covariates (except the intercept) were standardized, to have zero mean and variance 1. For the mean covariates, the original temporal derivative of the head motion parameters was used. For the variance covariates, the absolute value was used instead, as the variance should always increase at a motion spike regardless of the direction (positive or negative), see Figure 4 for an example of the co-variates for the mean and the variance. For both models, a fourth order AR model was used in each voxel. Variable selection was performed on all covariates (mean and variance), except for the intercept, as well as for the four AR parameters. Stationarity was enforced for the AR parameters, by discarding draws where the absolute value of any eigenvalue of the companion matrix is larger than or equal to 1. For each voxel, a total of 1,000 draws were used for MCMC burn-in and another 1,000 draws were saved for inference.

Figures 7-10 show receiver operating characteristic (ROC) curves for the two models, for different types (activity, motion, motion derivative, all) and levels (*γ* = 1,2 or 3) of heteroscedasticity. The ROC curves were generated by varying the threshold for the posterior probability maps (PPMs) from 0.01 to 1.00. It is clear that both models detect virtually all the active voxels for low levels of heteroscedasticity, while the homoscedastic model fails to detect a large portion of the active voxels with heteroscedastic noise for higher levels of heteroscedasticity. The posterior inclusion probabilities for the variance parameters (*γ*) indicate that the heteroscedastic model in virtually all voxels only includes the covariates that were used to generate the heteroscedastic noise (not shown).

**Figure 7:**
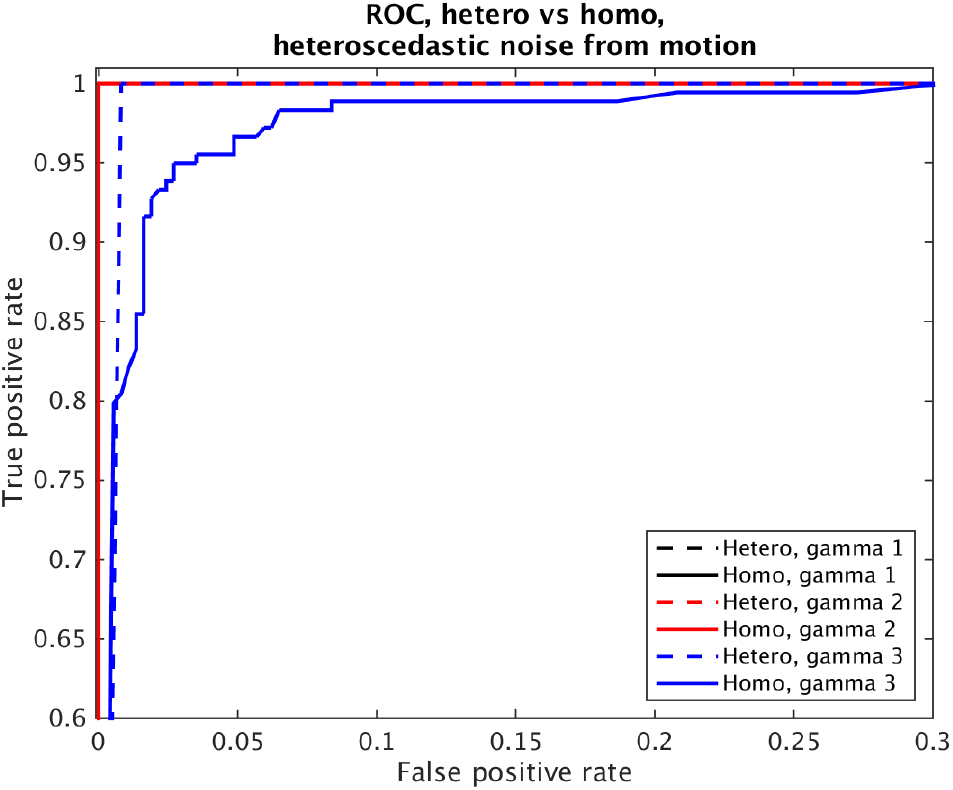
ROC curves for simulated data with heteroscedastic noise from one motion covariate, and different levels of heteroscedasticity. Both models perform well for low levels of heteroscedasticity, but the homoscedastic model performs worse for high levels of heteroscedasticity.

**Figure 8:**
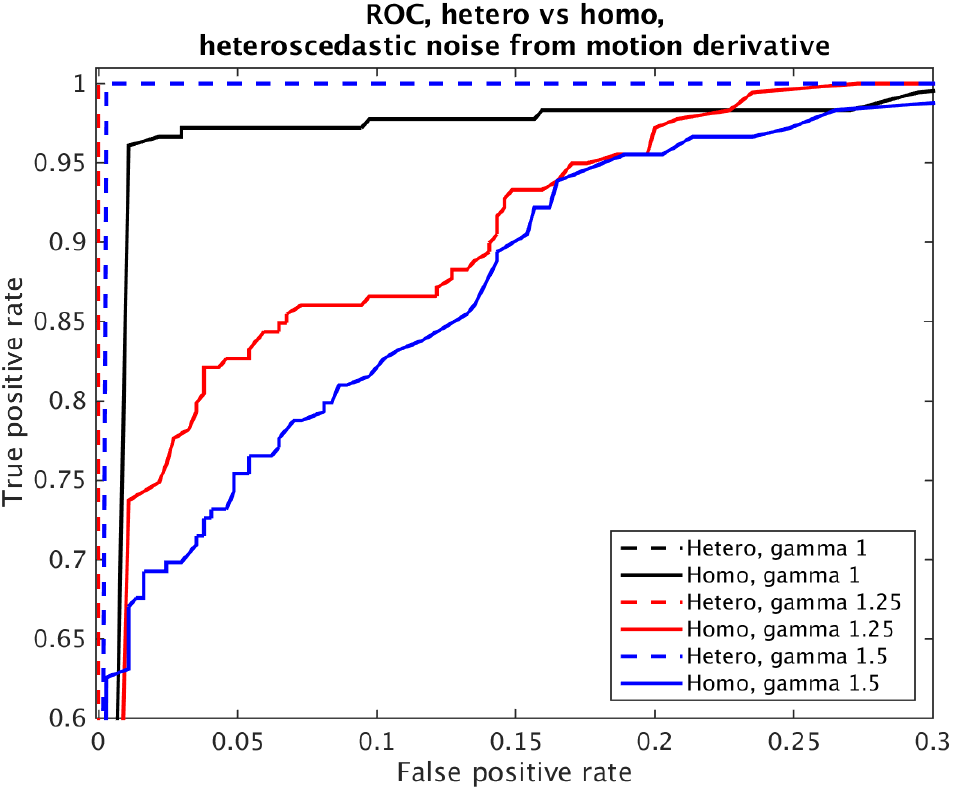
ROC curves for simulated data with heteroscedastic noise from the temporal derivative of one motion covariate, and different levels of heteroscedasticity. The homoscedastic model has a lower performance, and fails to detect a large portion of the active voxels.

**Figure 9:**
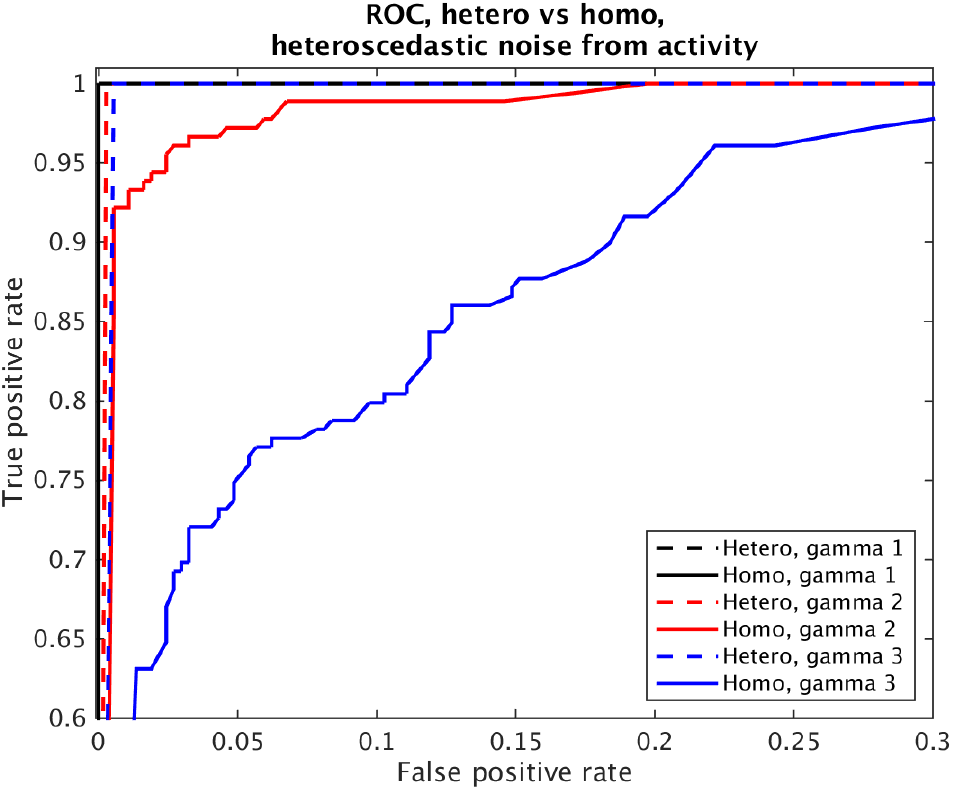
ROC curves for simulated data with heteroscedastic noise from one activity covariate, and different levels of heteroscedasticity. The homoscedastic model has a lower performance, and fails to detect a large portion of the active voxels.

**Figure 10:**
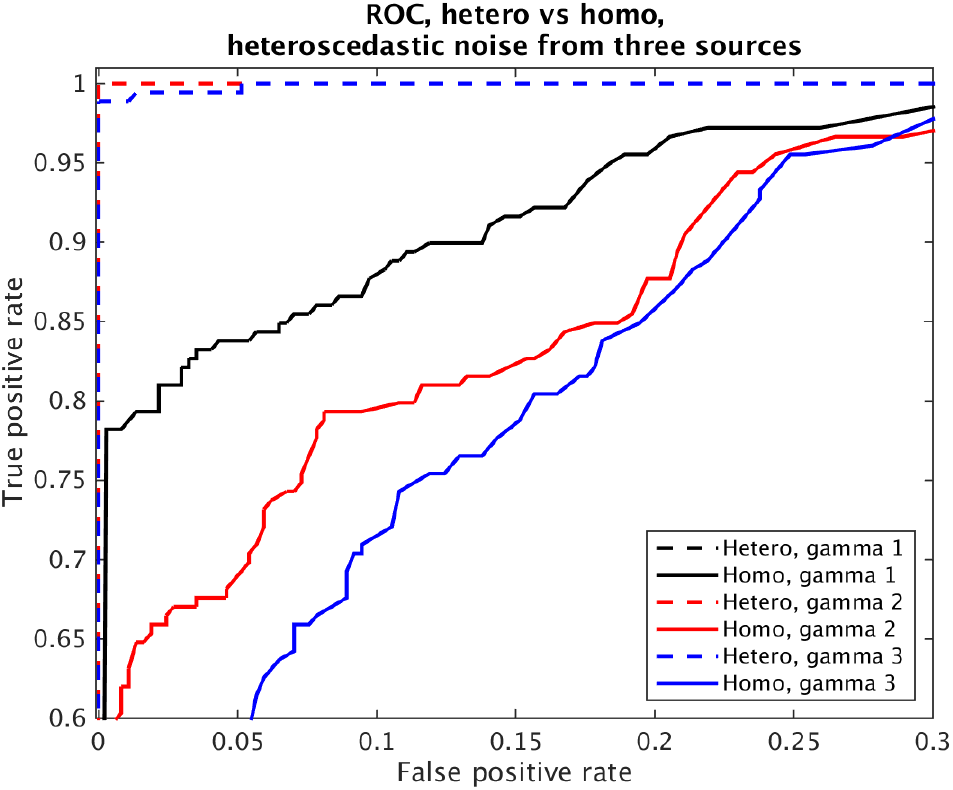
ROC curves for simulated data with heteroscedastic noise from three simultaneous sources (motion, motion derivative, activity), and different levels of heteroscedasticity. The homoscedastic model has a much lower performance, and fails to detect a large portion of the active voxels.

#### 5.1.2. GLMH vs weighted least squares

To compare the heteroscedastic model to the weighted least squares (WLS) approach proposed by Diedrichsen and Shadmehr [6], where a single weight is estimated for each volume, two additional datasets were simulated (using the same activity mask as above). For the first dataset, the same heteroscedastic noise was added to all voxels. For the second dataset, heteroscedastic noise was added to only 30% of the voxels (using the same hetero mask as above). The simulation was performed to generate different types (motion, motion derivative) and levels (*γ* = 1, 2 or 3) of heteroscedasticity. As the two approaches use different models for the temporal autocorrelation, the four AR parameters were set to 0, to focus solely on the heteroscedasticity. To mimic the analysis by Diedrichsen and Shadmehr [6], no motion regressors were used in the design matrix for the WLS approach. Bayesian *t*-scores (posterior mean / posterior standard deviation) were calculated for the heteroscedastic model, and compared to regular *t*- scores from the WLS approach. Figures 11-14 show ROC curves for the two approaches. Both approaches work well when the same heteroscedastic noise is present in all voxels, but the WLS approach fails to detect a large portion of the activity when the heteroscedastic noise is only present in 30% of the voxels.

**Figure 11:**
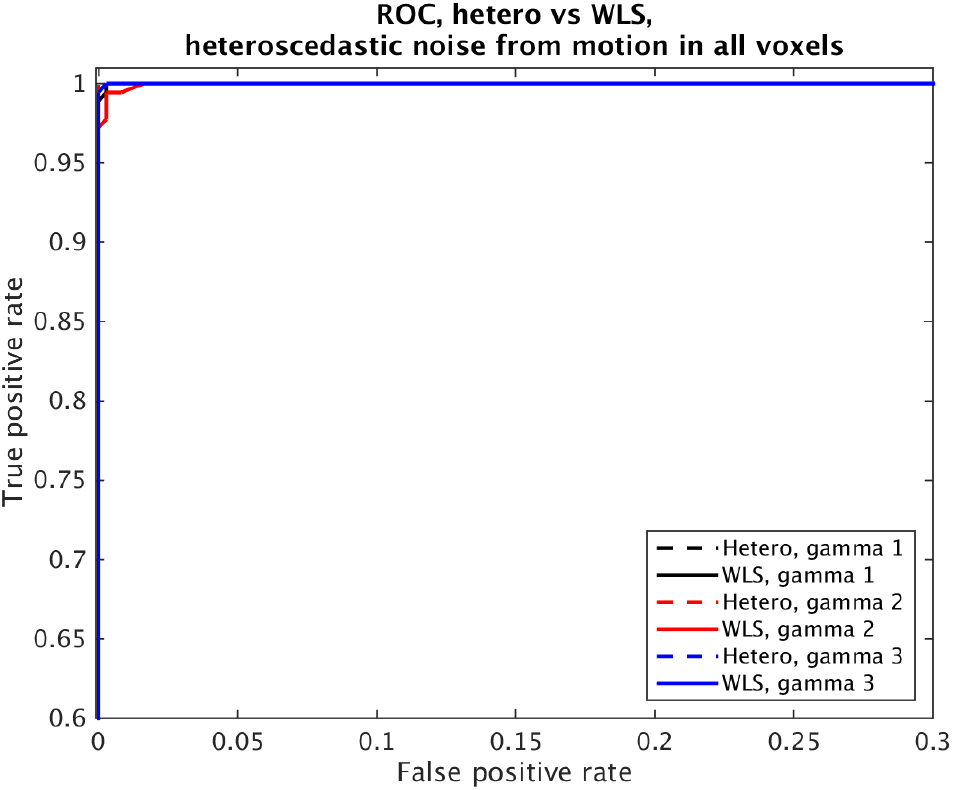
ROC curves for simulated data with heteroscedastic noise in all voxels, generated by one motion covariate. Both approaches perform well for all levels of heteroscedasticity.

**Figure 12:**
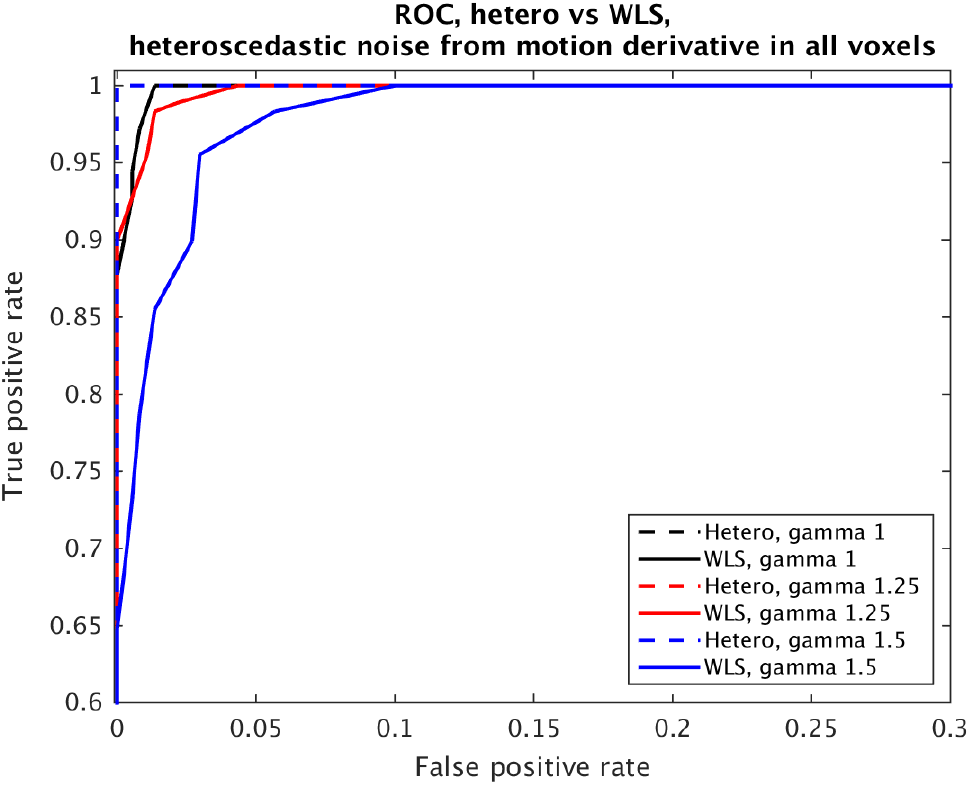
ROC curves for simulated data with heteroscedastic noise in all voxels, generated by the temporal derivative of one motion covariate. Both approaches perform well, but the hetero approach works better for higher levels of heteroscedasticity.

**Figure 13:**
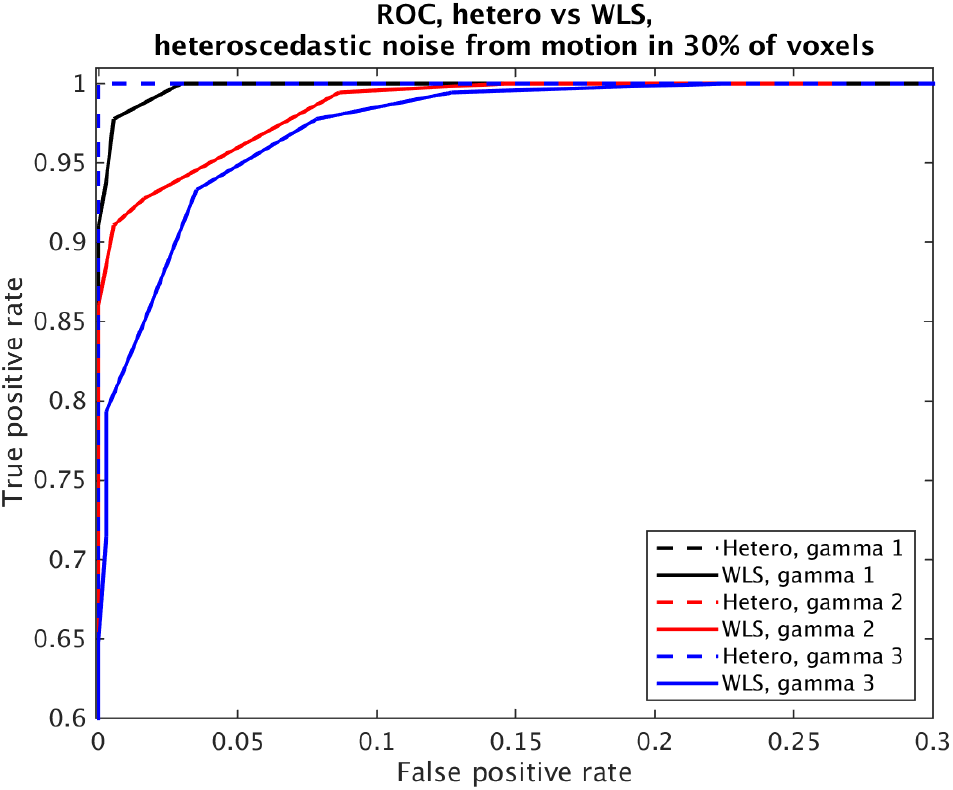
ROC curves for simulated data with heteroscedastic noise in 30% of the voxels, generated by one motion covariate. Compared to heteroscedastic noise in all voxels, the WLS approach has a slightly lower performance.

**Figure 14:**
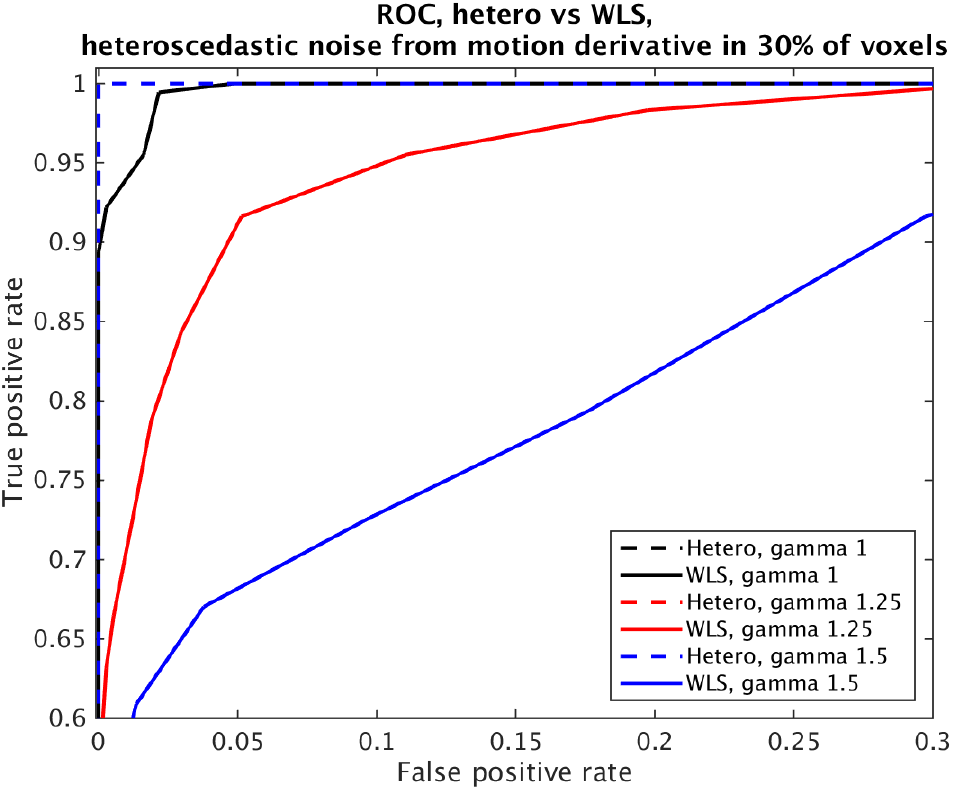
ROC curves for simulated data with heteroscedastic noise in 30% of the voxels, generated by the temporal derivative of one motion covariate. Compared to heteroscedastic noise in all voxels, the WLS approach fails to detect a large portion of the active voxels.

### 5.2. Application to real data

Three datasets from the OpenfMRI project[26, 27] were analyzed using both the homoscedastic and heteroscedastic noise models. The datasets include experiments on rhyme judgment1, living-nonliving judgment2 and mixed gambles3 [36].

In the rhyme judgment task, stimuli were presented in pairs (consisting of either words or pseudo-words) and the subject was asked whether the pair of stimuli rhymed with one another. The dataset consists of 13 subjects and two different conditions: words and pseudo-words.

In the living/nonliving judgment task, subjects were presented with words in either plain or mirror-reversed format, and asked whether the stimulus referred to a living or nonliving object. The data set consists of 14 subjects and 4 different conditions: mirror-reversed trials preceded by a plain text trial, mirror-reversed trials preceded by a mirror-reversed trial, plain-text trials preceded by a mirror-reversed trial, and plain-text trials preceded by a plain-text trial. A fifth covariate is used to represent failed (junk) trials.

Finally, in the mixed gambles task, subjects were presented with gambles in which they have a 50% chance of gaining and a 50% chance of losing money, where the potential gain and loss varied across trials. The subject then decided whether or not to accept the gamble. The data set consists of 16 subjects and 4 different conditions: task, parametric gain, parametric loss, and distance from indifference point. For more details on the 3 datasets we refer to the OpenfMRI website (https://openfmri.org).

#### 5.2.1. Single subject analysis

Prior to statistical analysis, the BROCCOLI software [9] was used to perform motion correction and 6 mm FWHM smoothing. For each subject, the analysis was performed as described for the simulated data (16 covariates + activity covariates, for both mean and variance). For each dataset, the analysis was performed (i) including only an intercept for the variance (i.e., a homoscedastic model) and (ii) including all covariates for the variance (i.e., a heteroscedastic model). Only gray matter voxels were analyzed to lower processing time. All results were finally transformed to MNI space, by combining T1-MNI and fMRI-T1 transforms.

Figure 15 shows PPMs for one subject from the rhyme judgment dataset and one subject from the mixed gambles dataset; the heteroscedastic model tends to detect more brain activity compared to the homoscedastic model. Figures 18 - 20 summarize the number of voxels where the difference between the heteroscedastic PPM and the homoscedastic PPM is larger than 0.5, for the three different datasets. The largest PPM differences are found in the rhyme judgment dataset, which contains the highest number of motion spikes. Figure 24 shows a comparison between the estimated homoscedastic and heteroscedastic standard deviation for a single time series; the heteroscedastic standard deviation is much higher for time points close to motion spikes, but lower for time points with little head motion. The homoscedastic model struggles to find a single variance to fit both time points with and without motion, thereby ending up inflating the variance at times with little or no motion. The heteroscedastic model can have a lower variance at timeperiods with little motion, and is therefore able to detect more brain activity. Figures 21 - 23 show the number of voxels, for each dataset, where the posterior inclusion probability is larger than 90% for the variance covariates. The temporal derivative of the head motion parameters are clearly the most important covariates for modeling the variance.

**Figure 15:**
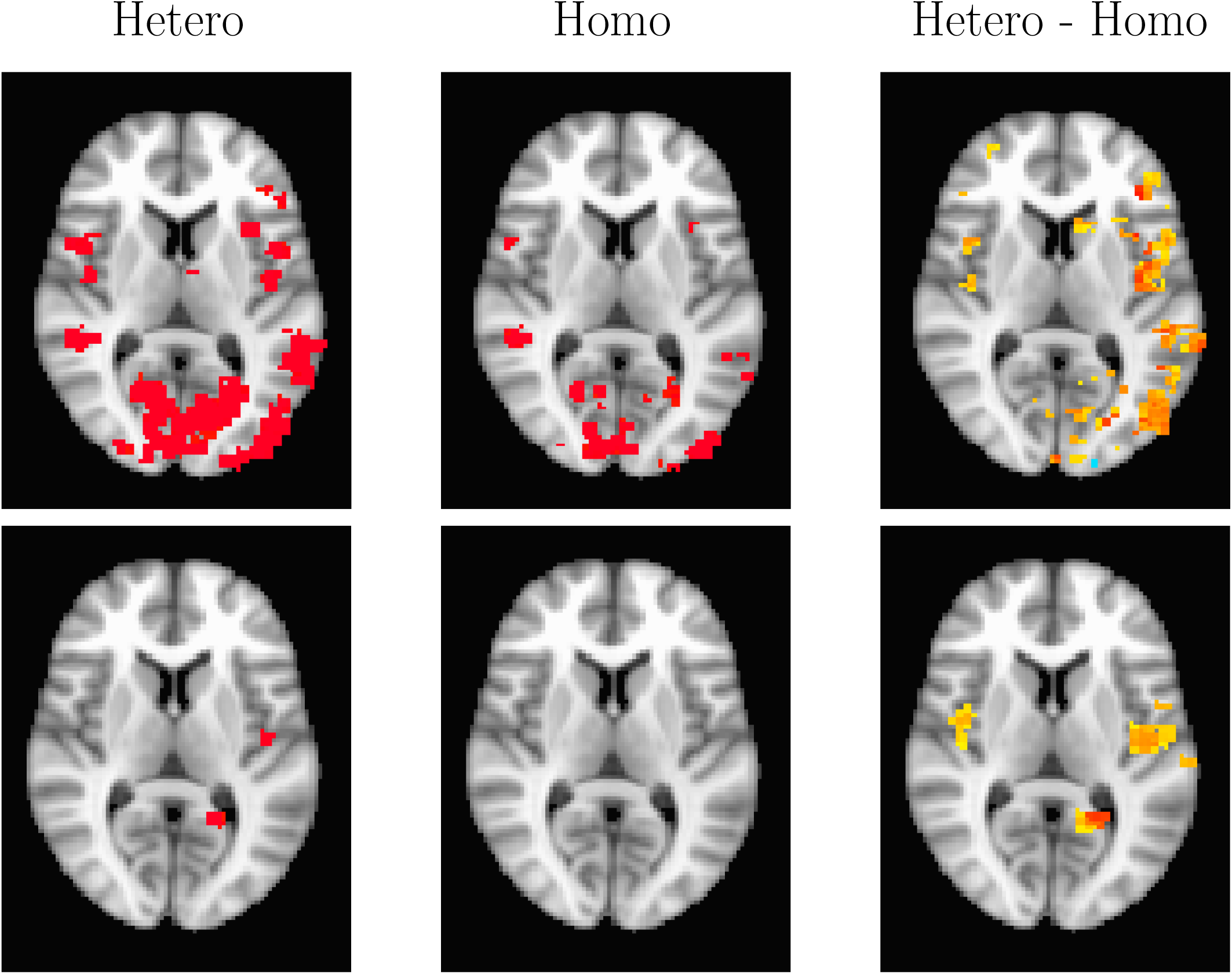
Single subject posterior probability maps (PPMs) for the rhyme judgment and mixed gambles datasets. From left to right: PPM for the heteroscedastic model, PPM for the homoscedastic model, PPM hetero - PPM homo. The hetero and the homo PPMs are thresholded at Pr = 0.95, while the difference is thresholded at 0.5. First row: Rhyme judgment dataset (subject 4, pseudo words contrast), Second row: Mixed gambles dataset (subject 3, parametric loss contrast). For subjects with one or several motion spikes, the heteroscedastic and the homoscedastic PPMs differ for a number of voxels. The reason for this is that the homoscedastic model overestimates the constant variance term, due to time points corresponding to motion spikes. The heteroscedastic model instead incorporates the head motion parameters, or the temporal derivative of them, to model these variance increases, and can thereby detect more brain activity.

#### 5.2.2. Sensitivity analysis

To investigate the importance of the prior settings, the analysis of the rhyme judgment dataset was repeated for the following prior settings.

Default: *τ_β_* = *τ_γ_* = 10, *τ_ρ_* = 1, *r* = 0.5, *ζ* =1,
Analysis 2: *τ_β_* = *τ_γ_* = 10, *τ_ρ_* = 0.5, *r* = 0.5, *ζ* =1,
Analysis 3: *τ_β_* = *τ_γ_* = 10, *τ_ρ_* = 1, *r* = 0.5, *ζ* = 0.5,
Analysis 4: *τ_β_* = *τ_γ_* = 10, *τ_ρ_* = 0.5, *r* = 0.5, *ζ* = 0.5,
Analysis 5: *τ_β_* = *τ_γ_* = 5, *τ_ρ_* = 1, *r* = 0.5, *ζ* =1,

Figure 16 shows the resulting homoscedastic and heteroscedastic PPMs for subject 4, which had the largest number of motion spikes. Lowering the prior variances *τ_β_* and *τ_γ_* leads to a clear decrease in detected brain activity, while the parameters for the noise process (*τ_ρ_*, *r*, and *ζ*) have a small effect on the detected brain activity.

**Figure 16:**
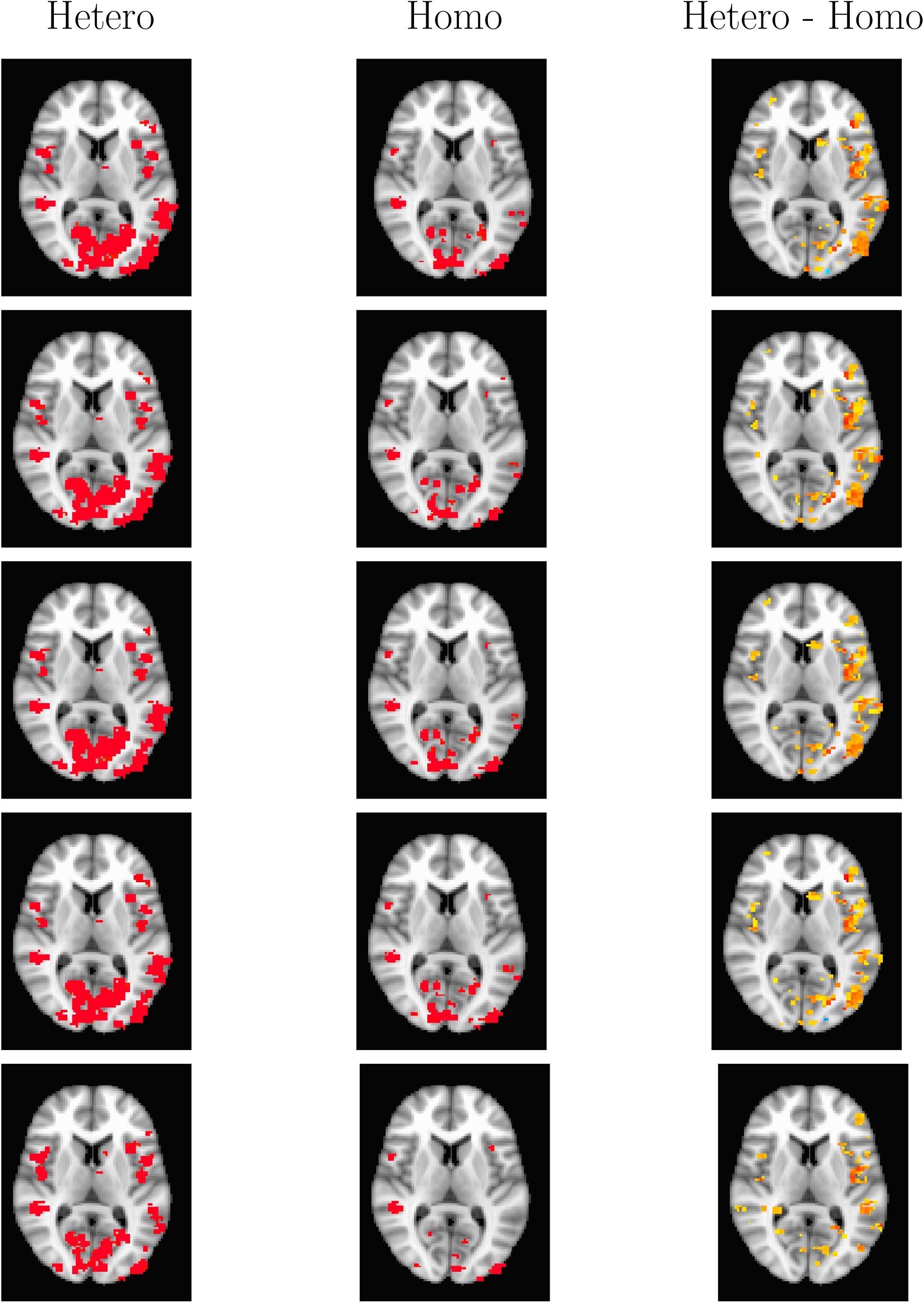
Single subject posterior probability maps (PPMs) for the rhyme judgment dataset (subject 4, pseudo words contrast). From left to right: PPM for the heteroscedastic model, PPM for the homoscedastic model, PPM hetero-PPM homo. The hetero and the homo PPMs are thresholded at Pr = 0.95, while the difference is thresholded at 0.5. First row: default prior parameters, *τ_β_* = *τ_γ_* = 10, *τ_ρ_* = 1, *r* = 0.5, *ζ* =1, Second row: *τ_β_* = *τ_γ_* = 10, *τ_ρ_* = 0.5, *r* = 0.5, *ζ* =1, Third row: *τ_β_* = *τ_γ_* = 10, *τ_ρ_* = 1, *r* = 0.5, *ζ* = 0.5, Fourth row: *τ_β_* = *τ_γ_* = 10, *τ_ρ_* = 0.5, *r* = 0.5, *ζ* = 0.5, Fifth row: *τ_β_* = *τ_γ_* = 5, *τ_ρ_* = 1, *r* = 0.5, *ζ* =1.

#### 5.2.3. Effect of updating π_β_ and π_γ_

To investigate the effect of updating *π_β_* and *π_γ_* in every MCMC iteration, compared to using fix values, the analysis of the rhyme judgment dataset was repeated with and without updating the inclusion parameters. Figure 17 shows the resulting homoscedastic and heteroscedastic PPMs for subject 4. Updating the inclusion parameters leads to lower posterior probabilities for the activity covariates, but the difference between the heteroscedastic and homoscedastic models is still rather large.

**Figure 17:**
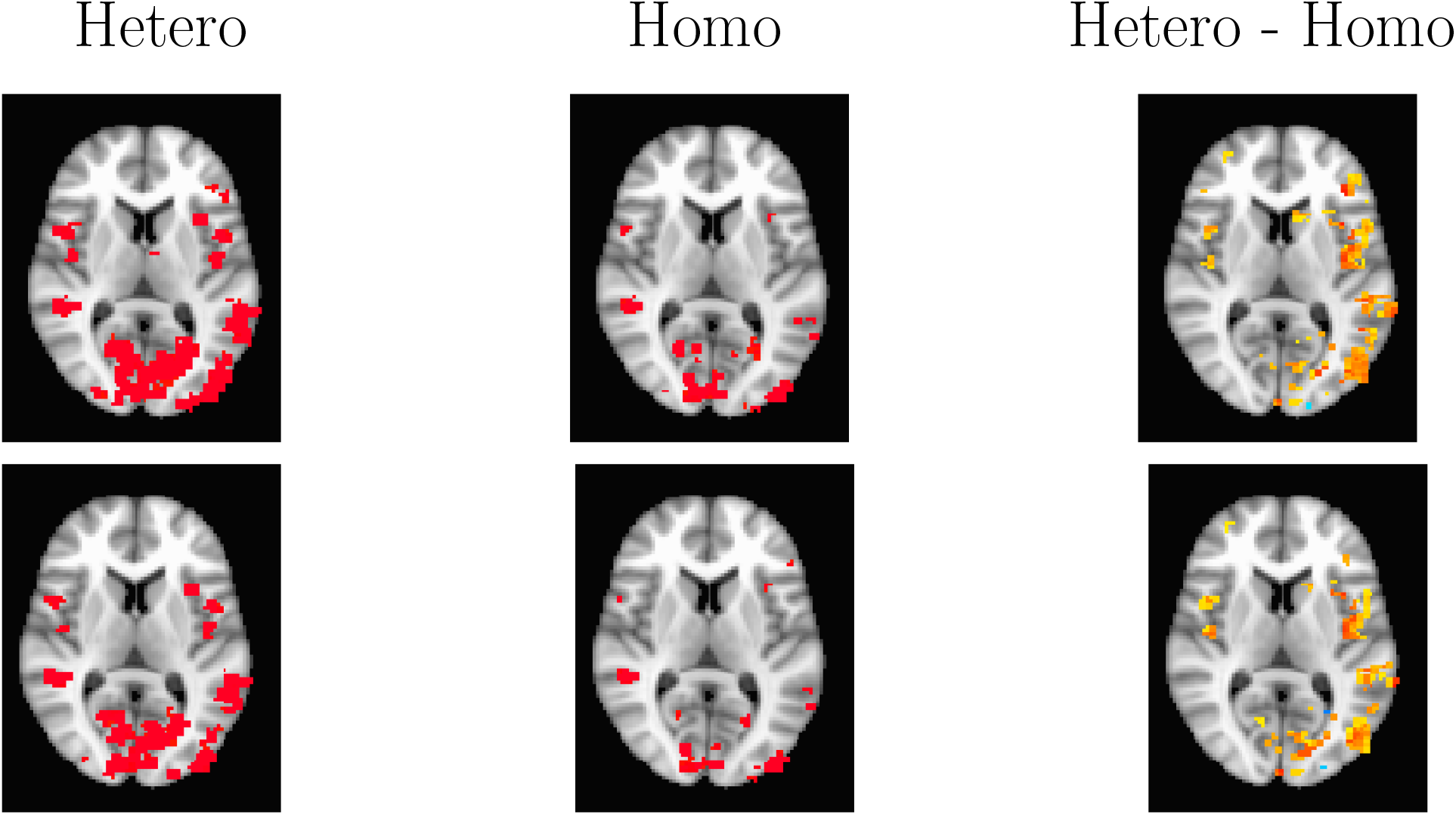
Single subject posterior probability maps (PPMs) for the rhyme judgment dataset (subject 4, pseudo words contrast). From left to right: PPM for the heteroscedastic model, PPM for the homoscedastic model, PPM hetero - PPM homo. The hetero and the homo PPMs are thresholded at Pr = 0.95, while the difference is thresholded at 0.5. First row: the inclusion parameters *π_β_* and *π_γ_* are fixated at 0.5, Second row: the inclusion parameters *π_β_* and *π_γ_* are updated in every MCMC iteration.

**Figure 18:**
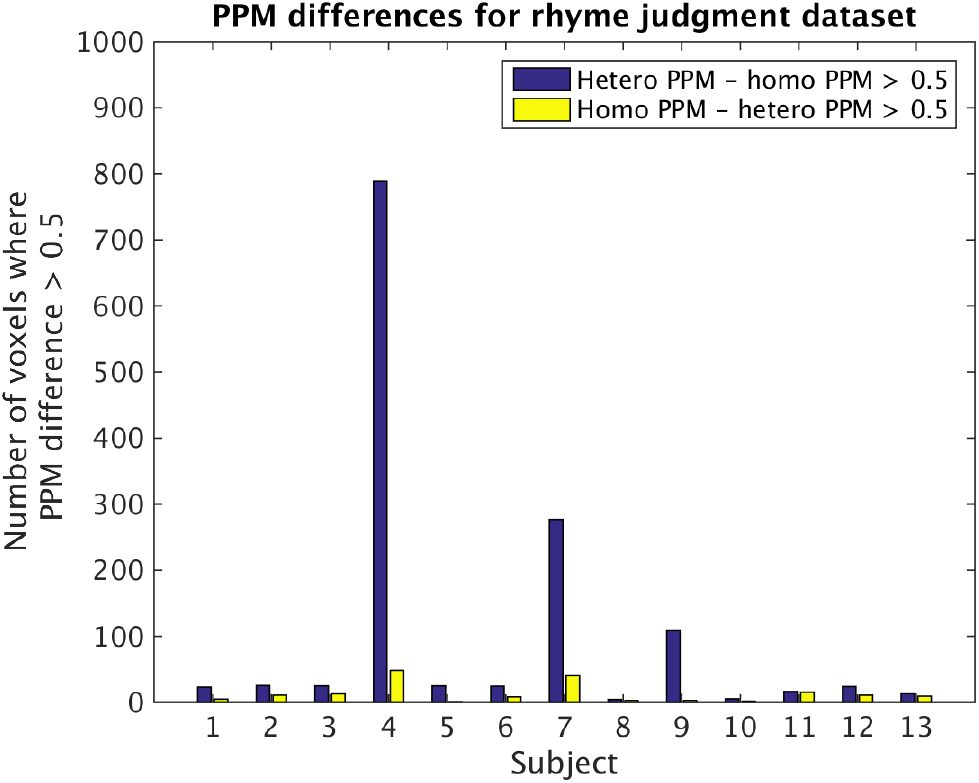
Number of gray matter voxels where the difference between the heteroscedastic PPM and homoscedastic PPM is larger than 0.5, for the rhyme judgment dataset.The bars represent the average over all activity covariates.

**Figure 19:**
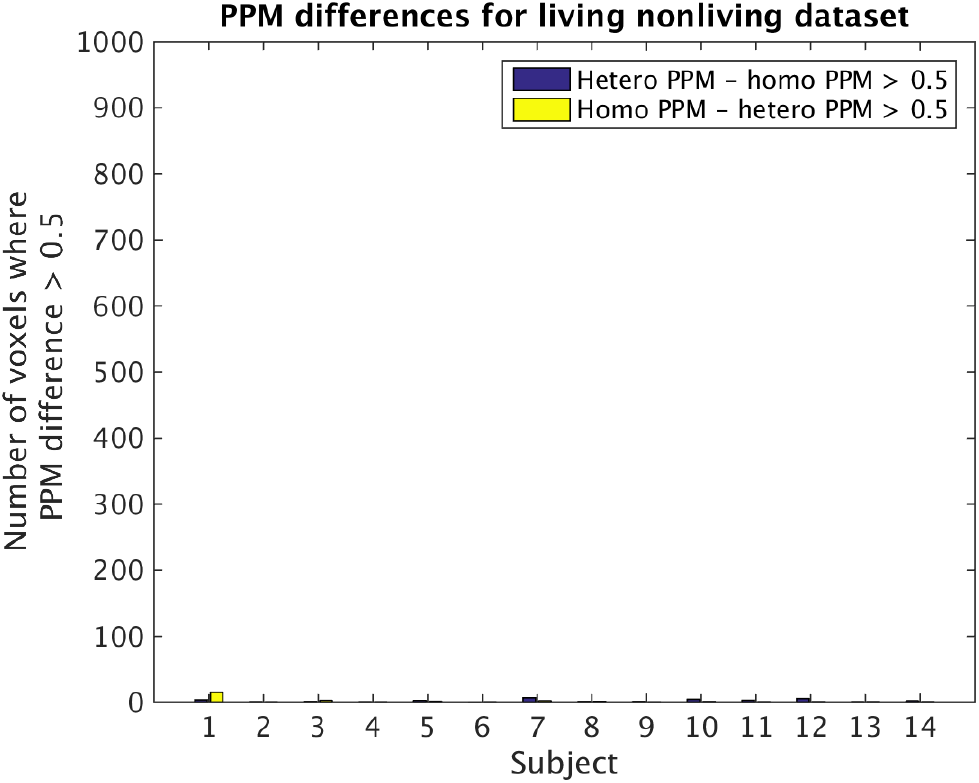
Number of gray matter voxels where the difference between the heteroscedastic PPM and homoscedastic PPM is larger than 0.5, for the living nonliving dataset. The bars represent the average over all activity covariates.

**Figure 20:**
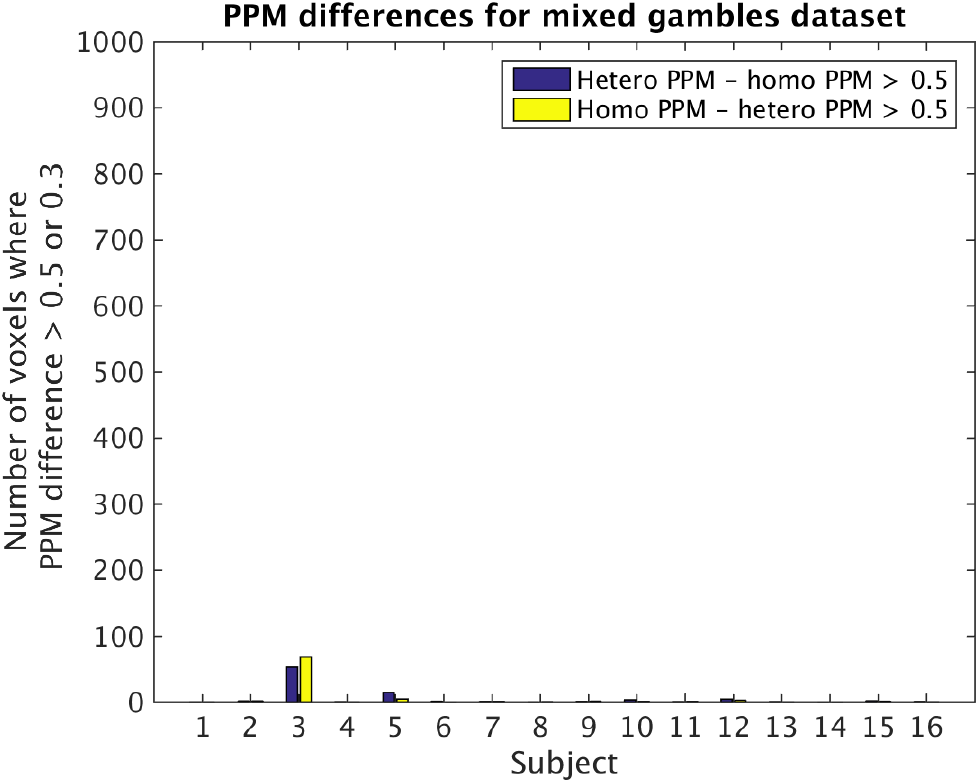
Number of gray matter voxels where the difference between the heteroscedastic PPM and homoscedastic PPM is larger than 0.5, for the mixed gambles dataset. The bars represent the average over all activity covariates.

**Figure 21:**
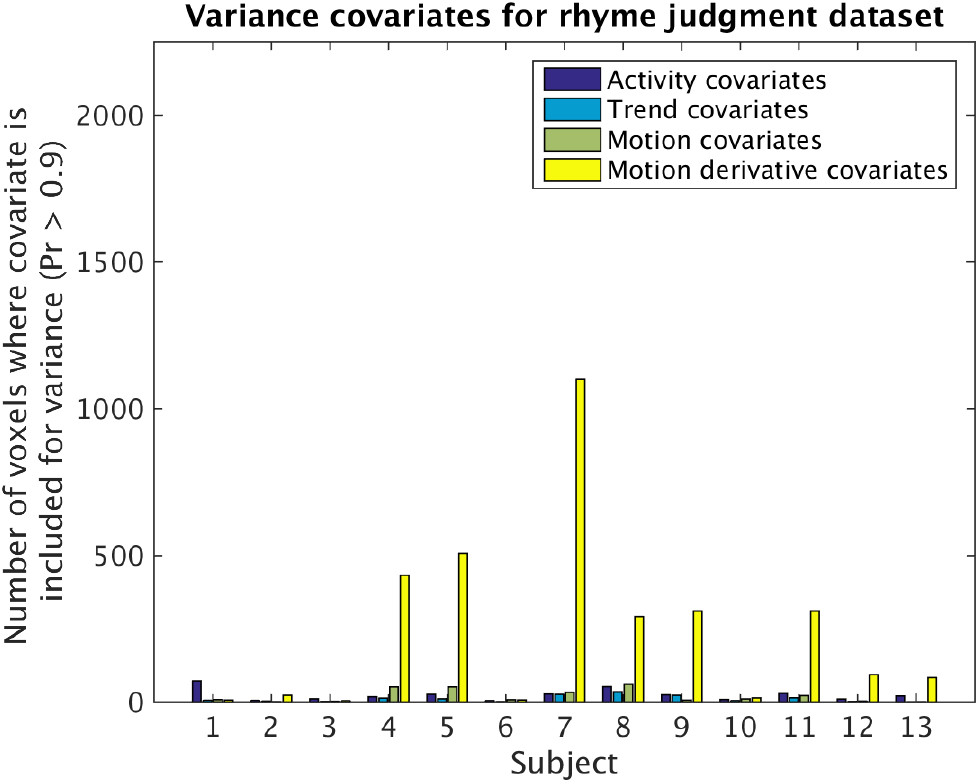
The use of variance covariates for the rhyme judgment dataset. Each bar represents the mean number of gray matter voxels, for each type of covariate (activity, trends, motion, motion derivative), for which the covariate is included to model the variance (posterior inclusion probability larger than 0.9). For subjects with motion spikes, one or several motion derivative covariates are used to model the heteroscedastic variance for a large number of voxels. The mean number of gray matter voxels is 15,600.

**Figure 22:**
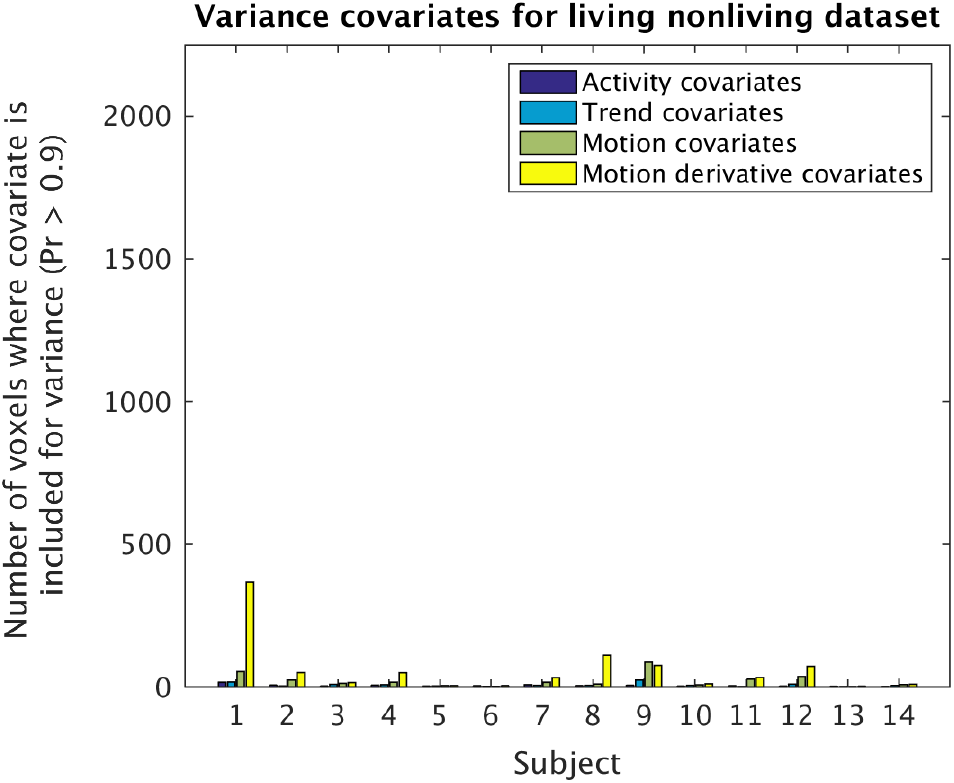
The use of variance covariates for the living nonliving dataset. Each bar represents the mean number of gray matter voxels, for each type of covariate (activity, trends, motion, motion derivative), for which the covariate is included to model the variance (posterior inclusion probability larger than 0.9). This dataset contains very few motion spikes, which explains why so few covariates are included in the variance. The mean number of gray matter voxels is 13,000.

**Figure 23:**
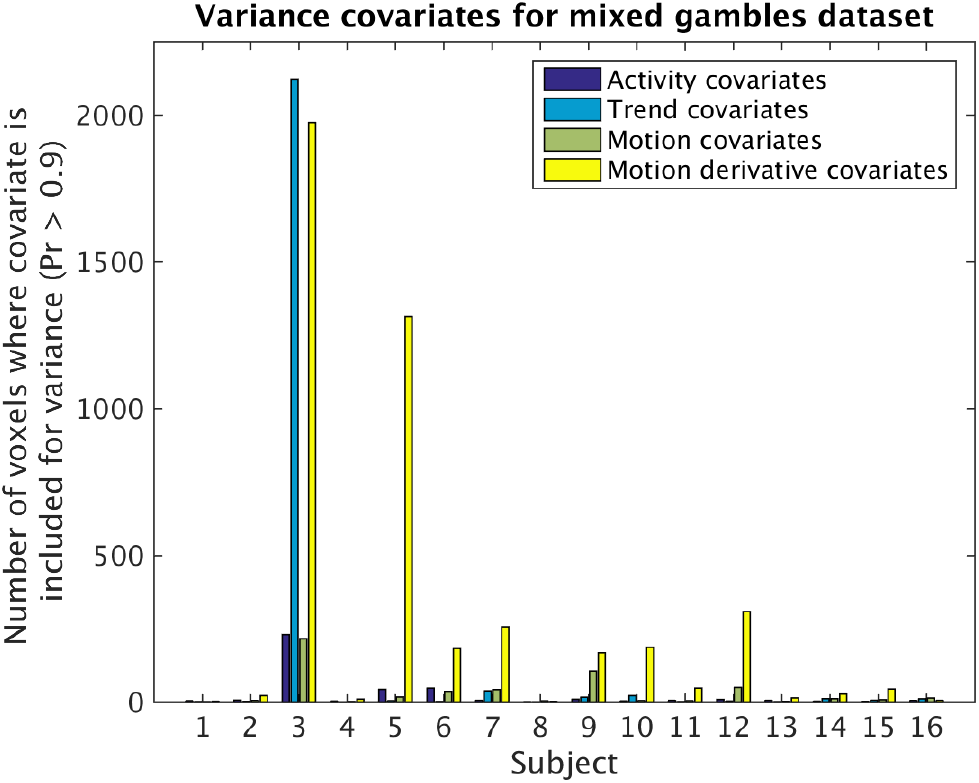
The use of variance covariates for the mixed gambles dataset. Each bar represents the mean number of gray matter voxels, for each type of covariate (activity, trends, motion, motion derivative), for which the covariate is included to model the variance (posterior inclusion probability larger than 0.9). For subjects with motion spikes, one or several motion derivative covariates are used to model the heteroscedastic variance for a large number of voxels. The mean number of gray matter voxels is 15,500.

**Figure 24:**
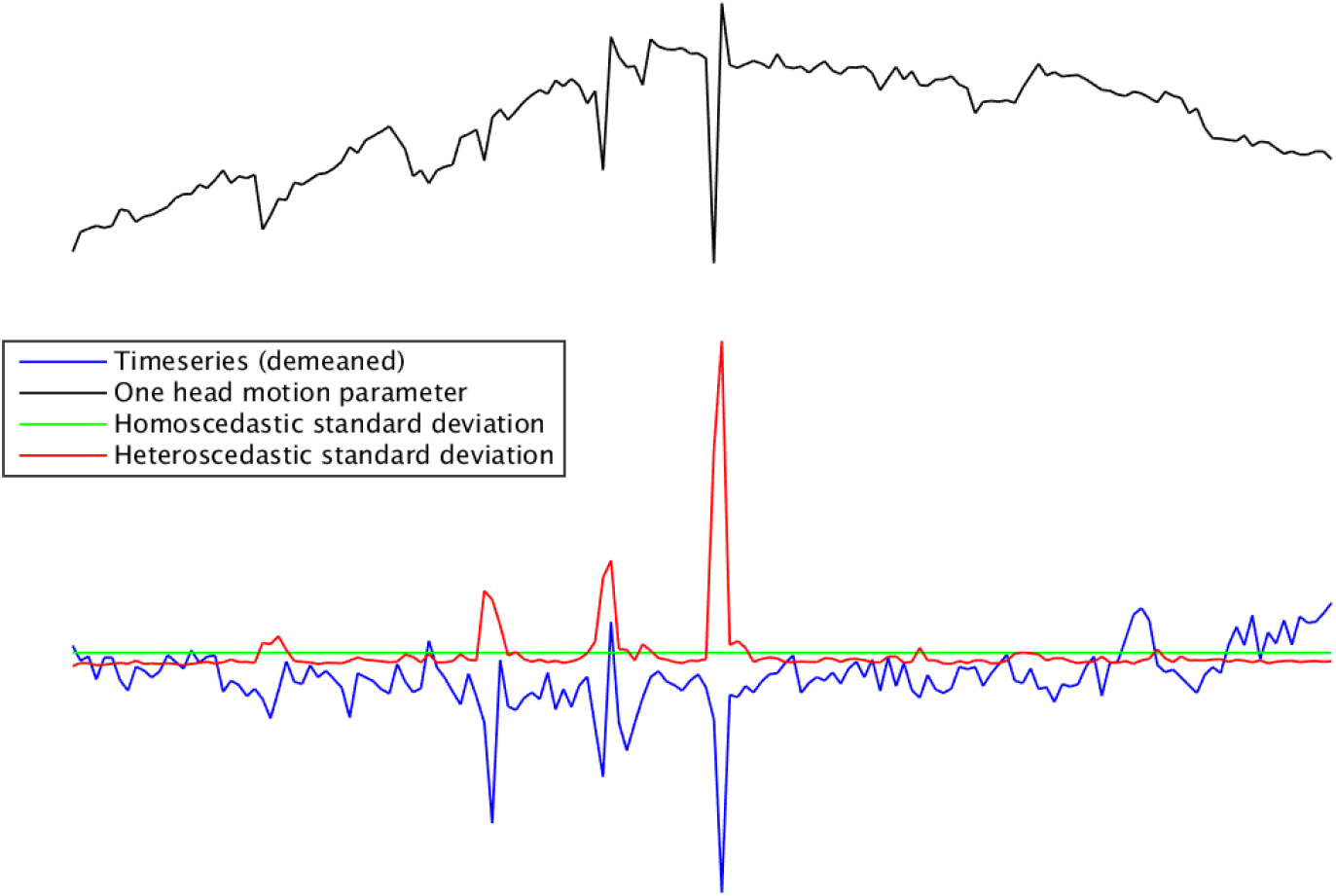
A comparison between the estimated homoscedastic and heteroscedastic standard deviation for one time series. The heteroscedastic standard deviation is much higher for the motion spikes, while it is lower for time points with little head motion. For this reason, the heteroscedastic model can automatically downweight time points close to motion spikes, and detect more brain activity by not over estimating the standard deviation for time points with little head motion.

#### 5.2.4. Convergence & efficiency of MCMC

The MCMC convergence is in general excellent; the acceptance probabilities for the variance covariates are 85.4% ± 5.1% for the rhyme judgment dataset, 89% ± 1.9% for the living nonliving dataset and 87% ± 7.1% for the mixed gambles dataset (standard deviation calculated over subjects). Trace plots are normally used to demonstrate convergence of MCMC chains, but the large number of voxels and covariates make such visual investigations difficult. For a single subject with 10,000 voxels in gray matter, the total number of trace plots would be 440,000 (representing

20 covariates for mean and variance and four AR parameters). The efficiency of the MCMC chain in each voxel was therefore instead investigated by calculating the inefficiency factor (also known as the integrated autocorrelation time [20] defined as 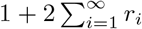, where *r_i_* is the ith autocorrelation of the MCMC draws for a given parameter) for each covariate for the mean and the variance, as well as for the four AR parameters. Since it is hard to estimate the inefficiency factor for variables with a low posterior inclusion probability (IPr), the inefficiency factor was only estimated if the IPr was larger than 0.3. To carefully investigate the MCMC efficiency in every voxel is difficult, due to the large number of voxels and covariates. An inefficiency factor of 1 is ideal, but very seldom achieved in practice. Inefficiency factors less than 10 - 20 are normally considered as acceptable. Tables 1, 2 and 3 therefore state the proportion of included voxels (IPr > 0.3) where the inefficiency factor is larger than 10, for the mean covariates (*β*), the variance covariates (*γ*), and the auto regressive parameters (*ρ*), respectively. The efficiency is in general high for both the mean and the variance covariates; only a few voxels have inefficiency factors larger than 10. The efficiency is in general lower for the auto regressive parameters, which has two explanations. First, the stationarity restriction enforces the parameters to a certain region, and if a parameter is repeatedly close to the boundary the sampling efficiency will be low. Second, in some voxels the algorithm finds a new mode after a subset of all the draws, which indicates that the chain has not converged. Considering the fact that 1,000 draws are already used for burnin, and that the processing time is 10 - 40 hours per subject, increasing the number of burnin draws even further is not a realistic option.

**Table 1:** Proportion of voxels with an inefficiency factor larger than 10 for the mean covariates (*β*), for the different datasets. The covariates in the design matrix have been grouped together to different types, and the numbers in the table represent the average over covariates (of each type) and subjects. The standard deviation was calculated over subjects.

| Dataset / Covariate type | Activity | Time trends | Motion parameters (MP) | Derivative of MP |
| --- | --- | --- | --- | --- |
| Rhyme judgment | $3.7\% \pm 1.9\%$ | $1.8\% \pm 1.0\%$ | $1.9\% \pm 0.9\%$ | $2.0\% \pm 1.1\%$ |
| Living nonliving decision | $2.0\% \pm 1.1\%$ | $1.7\% \pm 1.2\%$ | $1.5\% \pm 0.9\%$ | $2.2\% \pm 1.3\%$ |
| Mixed gambles task | $3.0\% \pm 1.8\%$ | $3.0\% \pm 1.9\%$ | $2.9\% \pm 1.7\%$ | $3.4\% \pm 2.1\%$ |

**Table 2:** Proportion of voxels with an inefficiency factor larger than 10 for the variance covariates (*γ*), for the different datasets. The covariates in the design matrix have been grouped together to different types, and the numbers in the table represent the average over covariates (of each type) and subjects. The standard deviation was calculated over subjects.

| Dataset / Covariate type | Activity | Time trends | Motion parameters (MP) | Derivative of MP |
| --- | --- | --- | --- | --- |
| Rhyme judgment | $1.3\% \pm 0.8\%$ | $1.0\% \pm 0.4\%$ | $1.0\% \pm 0.3\%$ | $0.8\% \pm 0.2\%$ |
| Living nonliving decision | $1.2\% \pm 0.5\%$ | $1.3\% \pm 0.5\%$ | $1.9\% \pm 0.5\%$ | $1.4\% \pm 0.3\%$ |
| Mixed gambles task | $1.8\% \pm 0.6\%$ | $1.8\% \pm 0.7\%$ | $2.1\% \pm 0.7\%$ | $1.7\% \pm 0.5\%$ |

**Table 3:** Proportion of voxels with an inefficiency factor larger than 10 for the auto correlation parameters (*ρ*), for the different datasets. The standard deviation was calculated over subjects.

| Dataset / AR parameter | AR 1 | AR 2 | AR 3 | AR 4 |
| --- | --- | --- | --- | --- |
| Rhyme judgment | $20.7\% \pm 3.0\%$ | $10.7\% \pm 2.4\%$ | $0.8\% \pm 0.7\%$ | $0.2\% \pm 0.3\%$ |
| Living nonliving decision | $12.2\% \pm 3.5\%$ | $10.3\% \pm 2.9\%$ | $1.2\% \pm 0.9\%$ | $0.1\% \pm 0.2\%$ |
| Mixed gambles task | $11.1\% \pm 3.5\%$ | $10.6\% \pm 2.7\%$ | $2.3\% \pm 1.8\%$ | $0.4\% \pm 0.6\%$ |

#### 5.2.5. Group analysis

Group analyses were performed using the full posterior of the task-related covariates from each subject (1,000 draws). To keep things simple, we perform each group analysis by computing the posterior for the sample mean: 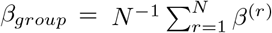, where *β*^(*r*)^ is the scalar activity coefficient for the *r*th subject in the sample, and *N* is the number of subjects. For each draw, the mean brain activity over subjects was calculated, to form the posterior of the mean group activity, *β*_group_. In a second group analysis, each subject was weighted with the inverse posterior standard deviation, i.e. 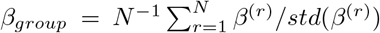. Figure 25 shows hetero and homo group mean PPMs (unweighted and weighted) for the rhyme judgment dataset, minimal differences were found for the other two datasets. The difference between the two models is slightly larger for the weighted group analysis, which is natural as the GLMH approach mainly affects the variance of the posterior. The effect of using a heteroscedastic model would clearly be stronger at the group level if many subjects (e.g. children) in the group exhibit motion spikes.

**Figure 25:**
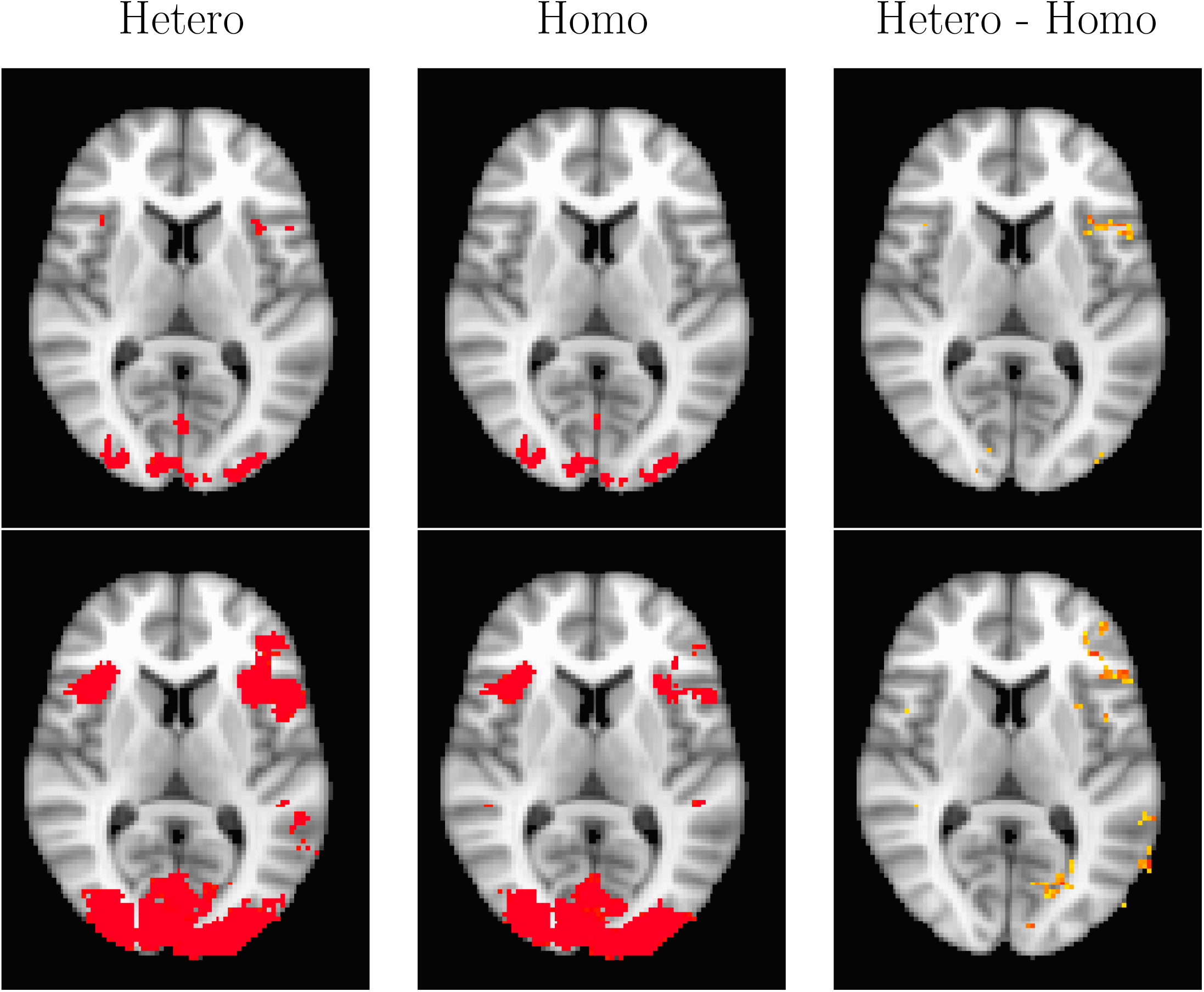
Group level posterior probability maps (PPMs) for the rhyme judgment dataset (contrast pseudo words). From left to right: PPM for the heteroscedastic model, PPM for the homoscedastic model, PPM hetero - PPM homo. The hetero and the homo PPMs are thresholded at Pr = 0.95, while the difference is thresholded at 0.5. Top row: group activity calculated without any subject specific weights. Bottom row: group activity calculated by weighting each subject with the inverse standard deviation.

## 6. Discussion

We have presented a Bayesian heteroscedastic GLM for single subject fMRI analysis. The heteroscedastic GLM takes into consideration the fact that the variance is inflated for time points with a high degree of head motion, and thus provides more sensitive results, compared to its homoscedastic counterpart. Instead of discarding data with too much head motion, or applying different scrubbing or censoring techniques [30, 28, 32], our heteroscedastic GLM automatically downweights the affected time points, and propagates the uncertainty to the group analysis by saving the full posterior. For the rhyme judgment dataset and the mixed gambles dataset, the temporal derivative of the head motion parameters are included as variance covariates for a large number of voxels. For heteroscedastic voxels in active brain areas, the difference between the homoscedastic PPM and the heteroscedastic PPM can be substantial. There will only be a sizeable PPM difference if the voxel belongs to an active brain area, and contains noise where the degree of heteroscedasticity is sufficiently high (see Figures 7 - 10). The difference between the two models is small for the living/nonliving dataset, mainly because that dataset contains very few motion spikes. This illustrates that our algorithm can be applied to any dataset, as using the heteroscedastic approach does not lead to a lower sensitivity when there are no motion spikes present.

### 6.1 MCMC vs Variational Bayes

A drawback of using MCMC is the computational complexity; it takes 10 - 40 hours (depending on the number of covariates) to analyze a single subject using the heteroscedastic model, with a single Intel Core i7 4790K CPU with 4 physical cores (8 cores due to hyper threading) and 32 GB of RAM. One alternative is to use variational Bayes (VB), where a few iterations is normally sufficient to obtain a point estimate of the posterior [24]. It is, however, much harder to perform variable selection within VB, and variable selection is necessary in our case since 18 - 21 covariates are used for the mean as well as for the variance. Without variable selection the model would contain too many parameters, compared to the number of time points in a typical fMRI dataset, which would result in poor estimates. Another problem with VB is that the posterior standard deviation is often underestimated.

In theory, the proposed algorithm can run on a graphics processing unit (GPU), which can analyze some 30,000 voxels in parallel [8, 9]. The pre-whitening step in each MCMC iteration is problematic from a GPU perspective, as a pre-whitened design matrix needs to be stored in each voxel / GPU thread. For 20 covariates and 200 time points, the design matrix requires 4,000 floats for storage. Modern Nvidia GPUs can, however, only store 255 floats per thread.

### 6.2. GLMH vs weighted least squares

To make a fair comparison between our heteroscedastic model and the WLS approach proposed by Diedrichsen and Shadmehr [6] is difficult, as we use Bayesian inference. Nevertheless, the WLS approach seems to work well as long as the same heteroscedastic noise is present in all voxels, but fails to detect activity when the heteroscedastic noise is only present in 30% of the voxels. Diedrichsen and Shadmehr [6] argue that the same weight should be used for all voxels, our results for real fMRI data (Figures 21-23) instead suggest that only a fraction of voxels have heteroscedastic noise. For some 13,000 - 15,600 voxels in gray matter, the derivative of the head motion parameters are included as covariates for the variance for 300 - 2,000 voxels (for subjects with motion spikes). Note that these numbers represent the average over each covariate type, meaning that if one of the six motion covariates is included for 12,000 voxels, the average over all six covariates will be 2,000 voxels.

The main drawback of the WLS approach is that it requires estimation of *T* weights from *T* time points, which results in extremely variable estimates unless the weights are averaged over many voxels. Our heteroscedastic GLM instead models the variance using a regression approach. Through the use of variable selection, a heteroscedastic model can be estimated independently in each voxel, even if the number of covariates is large.

### 6.3. Multiple comparisons

In contrast to frequentistic statistics, there is no consensus in the fMRI field regarding if and how to correct for multiple comparisons for PPMs. In this paper we have mainly focused on looking at differences between the heteroscedastic and the homoscedastic models, for voxel inference. It is not obvious how to use Bayesian techniques for cluster inference [10], which for frequentistic statistics has a higher statistical power. One possible approach is to use theory on excursion sets [3], to work with the joint PPM instead of marginal PPMs. Such an approach, however, requires a spatially dependent posterior, while we independently estimate one posterior for each voxel. One ad-hoc approach is to calculate a Bayesian t- or z-score for each voxel, and then apply existing frequentistic approaches for multiple comparison correction (e.g. Gaussian random field theory). This approach is for example used in the FSL software [39].

### 6.4. Future work

We have here only demonstrated the use of the heteroscedastic GLM for brain activity estimation, but it can also be used for estimating functional connectivity; for example by using a seed time series as a covariate in the design matrix. Although not investigated in this work, it is also possible to include additional covariates that may affect the variance, such as the global mean [29] or recordings of breathing and pulse [14]. Future work will also focus on adding a spatial model [25, 31], instead of analyzing each voxel independently.

## Acknowledgement

This work was financed by the Swedish Research council, grant 2013-5229 (“Statistical analysis of fMRI data”), and by the Information Technology for European Advancement (ITEA) 3 Project BENEFIT (better effectiveness and efficiency by measuring and modelling of interventional therapy). This research was also supported in part by NIH grants R01 EB016061 and P41 EB015909 from the National Institute of Biomedical Imaging. We thank Russ Poldrack and his colleagues for starting the OpenfMRI Project (supported by National Science Foundation Grant OCI-1131441) and all of the researchers who have shared their task-based data.

## 7. Appendix AImplementation

Our heteroscedastic GLM can be launched from a Linux terminal as

HeteroGLM fmri.nii.gz-designfiles activitycovariates.txt

gammacovariates gammacovariates.txt
ontrialbeta trialbeta.txt
ontrialgamma trialgamma.txt
ontrialrho trialrho.txt -mask mask.nii.gz
regressmotion motion.txt
regressmotionderiv motionderiv.txt
draws 1000 -burnin 1000 –savefullposterior
updateinclusionprob

where “activitycovariates.txt” states the activity covariates for the design matrix (normally only used for the mean), “gammacovariates.txt” states the covariates being used to model the variance, “ontrialbeta.txt” states covariates for which variable selection is performed for the mean, “ontrialgamma.txt” states covariates for which variable selection is performed for the variance and “ontrialrho.txt” states variable selection parameters for the autocorrelation. The option “updateinclusionprob” turns on updating the inclusion probabilities *π_β_* and *π_γ_* in every MCMC iteration. Covariates for intercept and time trends are automatically added internally. A homoscedastic GLM can easily be obtained as a special case, using only a single covariate (the intercept) for the variance. The following nifti files are created; posterior mean of beta and Ibeta (for each covariate), posterior mean of gamma and Igamma (for each covariate), posterior mean of rho and Irho (for each AR parameter), and PPMs for each activity covariate. The full posterior of all beta, gamma and rho parameters can also be saved as nifti files.

## 8. Appendix B - MCMC Details

Variable selection by MCMC in the linear regression model Let us assume a general multivariate prior *β_ℐ_\ℐ ~ N* (***μ***, Ω_*I*_). Now,

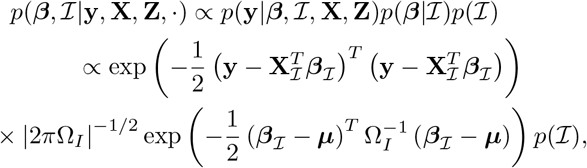

where **X**_ℐ_ is the matrix formed by selecting the columns of **X** corresponding to ℐ. The conditional likelihood 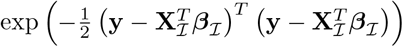 can be decomposed as

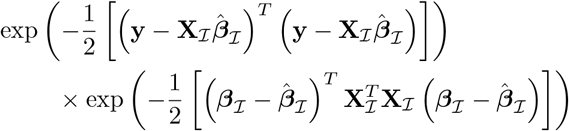

Multiplying the conditional likelihood by the prior and completing the square 4 gives

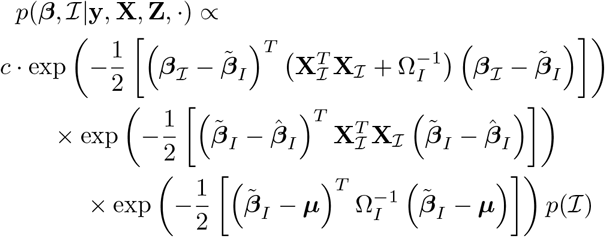

where 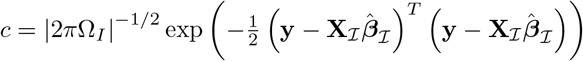 and 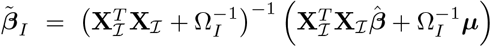 This shows that

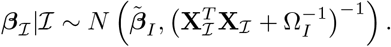

Integrating with respect to *β_ℐ_* gives

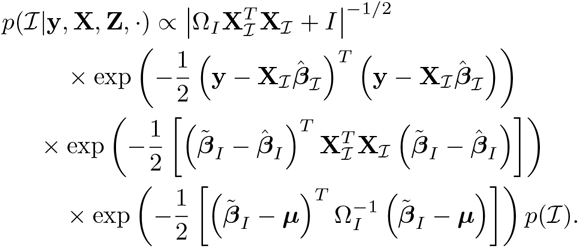

## Footnotes

1 https://openfmri.org/dataset/ds000003/

2 https://openfmri.org/dataset/ds000006/

3 https://openfmri.org/dataset/ds000005/

4 (*x-a*)*′A*(*x-a*)+(*x-b*)*′B*(*x-b*) = (*x-d*)*′D*(*x-d*)+(*d-a*)*′A*(*d-a*) + (*d - b*)*′B*(*d - b*), where *D = A + B* and *d* = *D*^-1^*(Aa + Bb):*

